# Dynamic Modulation of Representational Trajectories Through Selective Attention

**DOI:** 10.1101/2024.01.21.576506

**Authors:** Yu Zhou, Liang Zhang, Nikolai Axmacher, Daniel Pacheco Estefan, DaHui Wang, Yujian Dai, Xiaojing Peng, Gui Xue

## Abstract

Mounting evidence indicates that information processing in the visual hierarchy follows a sequential progression from low-level perceptual to high-level conceptual features during visual perception, with the reverse order during memory retrieval. However, the nature of this hierarchical processing and its modulation by selective attention remains unclear. By integrating a deep neural network with the drift-diffusion model on reaction time data, we identified parallel processing of perceptual and conceptual features during perception, rather than sequential processing. The slower reaction times observed in conceptual tasks compared to perceptual tasks were primarily driven by the larger decision boundary and longer non-decision time associated with conceptual features, which could not be compensated by their faster evidence accumulation rates. Using single-trial multivariate decoding of MEG (magnetoencephalography) data to examine the timing of neural representations of different features, we found that selective attention reversed the onset times of perceptual and conceptual features in the occipital and parietal lobes during perception. This led to earlier detection of conceptual features compared to perceptual features in the animacy and size tasks. During retrieval, the colour task exhibited earlier peak times for perceptual features compared to conceptual features in the frontal lobe, indicating that perceptual features were reconstructed with the highest fidelity before conceptual features. These findings provide novel insights into the hierarchical nature of information processing during perception and retrieval, emphasizing the crucial role of selective attention in modulating the speed of feature information accumulation.

## Introduction

Visual object recognition relies on a hierarchical information processing pathway in the ventral visual stream, progressing from low-level perceptual details to higher-level conceptual features (Carlson et al., 2013; Cichy et al., 2014; Khaligh-Razavi et al., 2018; Wang et al., 2022). During object recognition, low-level perceptual features were discriminated more quickly and could be decoded from brain activity earlier than high-level conceptual features (Clarke et al., 2013; Linde-Domingo et al., 2019; Mirjalili et al., 2021). Conversely, during memory retrieval, this sequence was reversed. Reaction times and brain activity patterns indicated that conceptual information was reconstructed more rapidly than perceptual details (Linde-Domingo et al., 2019; Mirjalili et al., 2021).

Two major perspectives on the timing of object categorization have been proposed to explain this phenomenon. One view suggests that the relative timing reflects a sequence of stages in visual processing linked to specific levels of object categorization, such as categorizing objects based on low-level perceptual features or high-level conceptual features. According to this perspective, fast categorizations are fast because they precede other categorizations within the visual processing hierarchy (Brincat & Connor, 2004; Serre et al., 2007; Yau et al., 2013). Another perspective posits that the relative timing reflects the amount of evidence required for the decision and the speed of evidence accumulation: fast indicates narrow boundary separation and/or a shorter time to accumulate the required amount of evidence used to drive a decision process (Ratcliff, 1978; Ratcliff & McKoon, 2008a).

One potential way to adjudicate between these alternative perspectives is to explore how selective attention modulates the order of these processes during encoding and retrieval. Selective attention prioritizes neural representations most relevant to current behavioural goals (Moore & Zirnsak, 2017). Mounting evidence indicated that selective attention could enhance target information and suppress non-target information. For example, studies showed that selective attention could enhance the activation of brain regions responsible for target processing (Baldauf & Desimone, 2014; Maunsell & Treue, 2006). Beyond activation strength, studies employing multivariate pattern analyses (MVPA) (Hebart & Baker, 2018; Kriegeskorte et al., 2006) have further found that selective attention could enhance the strength and fidelity of target representations relative to non-targets (Grootswagers et al., 2021; Long & Kuhl, 2018). Furthermore, attention has been shown to increase the representational dimension (Sheng et al., 2022) and the representational distance among target items, allowing for finer discrimination (Nastase et al., 2017).

Based on these findings, we hypothesize that if the processing hierarchy reflects the order or sequence of information processing, selective attention could modulate the relative time differences between different features but not the processing order. In contrast, if these features are processed in parallel and the speed reflects the amount and/or the speed of evidence accumulation, selective attention could potentially reverse the order of information processing. This would result in the intriguing possibility of detecting conceptual features earlier than perceptual features during object perception. We further compared the effect of selective attention on visual perception and memory retrieval, as existing studies have revealed that these two distinct processes exhibited reversed timing for perceptual and conceptual features (Linde-Domingo et al., 2019; Mirjalili et al., 2021).

To investigate this, we manipulated both perceptual (colour) and conceptual features (animacy and real-world size) of objects in a magnetoencephalography (MEG) experiment. Two tasks were conducted: a feature recall task and a feature perception task. In the feature recall task, participants learned object-location associations and subsequently recalled the target features of the cued objects. In the feature perception task, participants categorized objects based on the target features. Using single-trial decoding, we found selective attention reversed the order of onset times of perceptual and conceptual features in the occipital and parietal regions during perception, and the order of peak times in the frontal lobe during retrieval. Our findings provide new insights into the nature of processing hierarchy and reveal a novel mechanism of selective attention in modulating the temporal dynamics of information processing.

## Results

### Behavioural performance

We used three tasks, colour, animacy, and real-world size, to examine how task requirements modulate feature processing during perceptual and retrieval stages. The feature retrieval task preceded the feature perception task to mitigate potential influences of stimulus familiarity on memory retrieval. For the feature retrieval task, participants first learned the picture-location associations. Task cues were presented at the beginning of each memory block, prompting participants to retrieve specific features (colour, animacy, or real-world size) of the objects indicated by a particular location (Figure 1A). To further probe the verbatim memory, a recognition task followed, wherein participants were tasked with selecting the target items from a set of three similar pictures. For the feature perception task, participants were asked to categorize the object based on a specific feature (Figure 1B. also see Methods).

**Figure 1.**
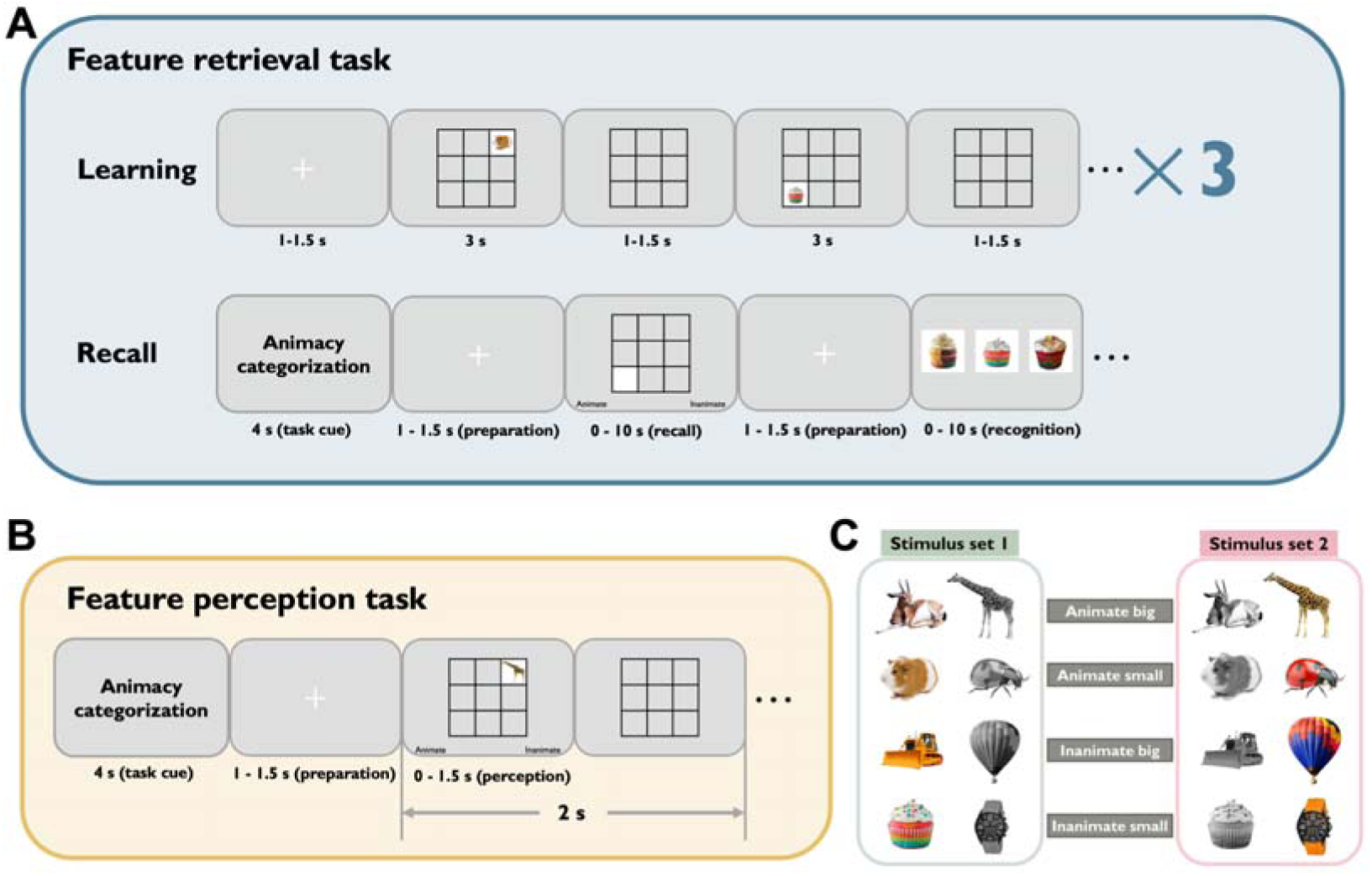
Experimental design and stimuli examples. **A. Feature retrieval task.** Participants first learned object-location associations and then recalled the specific features of cued objects, such as colour, animacy, or size. The figure shows animacy as an example. Participants were then tested on their ability to recognize the target objects they had seen during the learning phase. **B. Feature perception task.** In this task, participants categorized objects based on the target feature indicated during the task cue period. When presented with the objects, participants were tasked with categorizing them as quickly and accurately as possible. For example, if the task cue was ‘Animacy categorization’, participants were required to categorize the presented object as either animate or inanimate. **C. Object dimensions.** A total of 192 objects varied along three dimensions: one perceptual dimension (colorful or grayscale), and two conceptual dimensions: animacy (animate or inanimate) and real-world size (small or big).

In the first analysis, we aimed to examine the processing stream of different features through reaction time. Based on previous findings (Clarke et al., 2013; Linde-Domingo et al., 2019; Mirjalili et al., 2021), we hypothesized that during the visual perception stage, reaction times for perceptual features would be faster than those for conceptual features. Only correct trials were used for all reaction time (RT) analyses. Our results align with previous research. In the feature perception task, the mean reaction times were 0.699 s (*SD* = 0.092 s) for the colour task, 0.770 s (*SD* = 0.104 s) for the animacy task, and 0.798 s (*SD* = 0.088 s) for the size task. We used a linear mixed-effects model to directly test the differences in reaction times across the three tasks. The task type (colour, animacy, and size) was used to predict reaction times, with participant IDs included as a random effect. Task type significantly predicted reaction times (*F_2, 52_* = 48.738, *p* < 0.001). Post-hoc tests indicated that the reaction time for the colour task was faster compared to both the animacy (*t_52_* = -6.876, *p_corrected_*< 0.001) and size tasks (*t_52_* = -9.574, *p_corrected_* < 0.001), and reaction time in the animacy task was faster compared to the size task (*t_52_* = -2.698, *p_corrected_* = 0.025) (Figure 2A). In the feature retrieval task, we did not observe reversed patterns for reaction times compared to the feature perception task. The mean reaction times were 2.083 s (*SD* = 0.837 s) for the colour task, 2.048 s (*SD* = 0.682 s) for the animacy task, and 2.508 s (*SD* = 0.719 s) for the size task. The fixed factor task type significantly predicted reaction times (*F_2, 52_* = 16.124, *p* < 0.001). Post-hoc testing revealed that the reaction time for the size task was significantly longer compared to both the colour and animacy tasks (size task vs colour task: *t_52_* = 4.713, *p_corrected_* < 0.001; size task vs animacy task: *t_52_* = 5.100, *p_corrected_* < 0.001) (Figure 2B).

**Figure 2.**
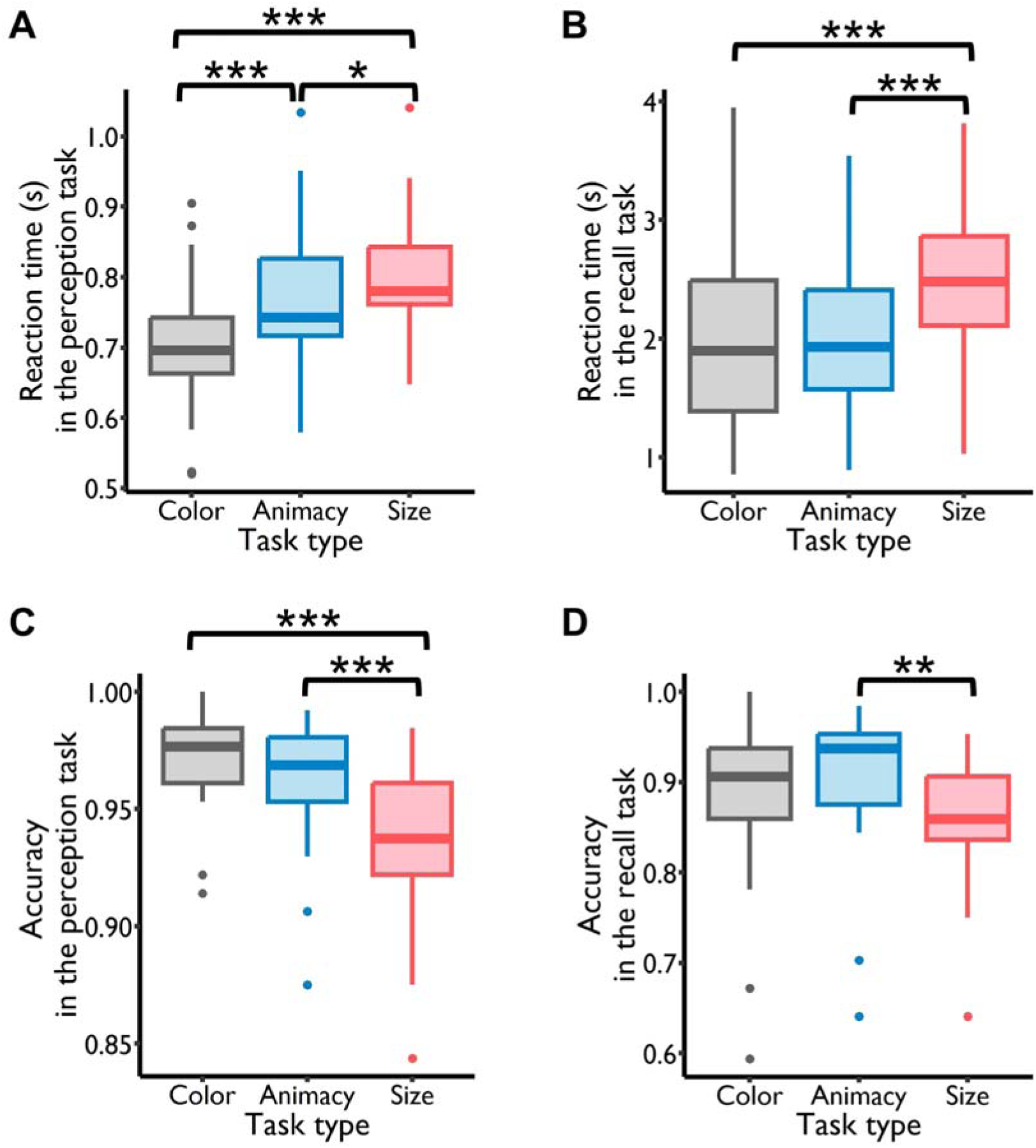
Behavioural results during the perception and retrieval tasks. **A.** Analysis of reaction times (RTs) for the colour, animacy and size perception tasks showed a significantly increasing pattern: RT_(colour)_ < RT_(animacy)_ < RT_(size)_, with all comparisons being statistically significant. **B.** Reaction times in the size retrieval task were significantly longer than in both the colour and the animacy retrieval tasks. **C.** Accuracy in both colour and animacy perception tasks significantly surpassed accuracy in the size task. **D.** Accuracy in the animacy retrieval task was significantly higher than in the size retrieval task. *: *p* < 0.05, **: *p* < 0.01, ***: *p* < 0.001.

Next, we assessed the accuracy in correctly perceiving or retrieving the features in each task type. The accuracy in the feature perception task was close to the ceiling for the colour and animacy tasks (colour task: *Mean* = 97.05%, *SD* = 2.24%; animacy task: *Mean* = 96.30%, *SD* = 2.69%), and high for the size task (size task: *Mean* = 93.58%, *SD* = 3.15%). The fixed factor task type significantly predicted participants’ accuracy (*F_2, 52_* = 24.435, *p* < 0.001). Post-hoc tests indicated that both the colour and animacy tasks showed significantly higher accuracy than the size task (colour task vs size task: *t_52_* = 6.644, *p_corrected_* < 0.001; animacy task vs size task: *t_52_* = 5.205, *p_corrected_* < 0.001). The accuracy of the colour and animacy tasks were comparable (*t_52_* = 1.440, *p_corrected_* = 0.328) (Figure 2C). In the feature retrieval task, the mean accuracy was 89.06% (*SD* = 9.37%) for the colour task, 90.39% (*SD* = 7.99%) for the animacy task, and 86.23% (*SD* = 7.02%) for the size task. The fixed factor task type significantly predicted participants’ accuracy (*F_2, 52_*= 5.913, *p* < 0.01). Post-hoc testing revealed that the accuracy of the animacy task was significantly higher compared to that of the size task (animacy task vs size task: *t_52_* = 3.366, *p_corrected_* < 0.01), while the accuracy of the colour task did not exhibit a significant difference compared to the other two tasks (colour task vs animacy task: *t_52_* = -1.075, *p_corrected_*= 0.533; colour task vs size task: *t_52_* = 2.291, *p_corrected_* = 0.066) (Figure 2D). The behavioural results of the recognition task are presented in Figure S1.

### Stimulus features are represented in parallel in the deep neural network

To explore whether the progressively slower reaction times for the colour, animacy, and size features were attributable to their positions at different stages within the visual processing hierarchy, we utilized the deep neural network (DNN) AlexNet, which mirrors the hierarchical structure of neurons along the ventral visual stream (Krizhevsky et al., 2017). AlexNet comprises eight layers, including five convolutional layers and three fully connected layers. Broadly, the early layers of AlexNet process early visual features such as colours, contrasts, and frequencies. In contrast, the deeper layers are responsible for higher-order visual representations, such as the surface structure of objects or body parts of animals. We proceeded with the assumption that if the layers in AlexNet containing colour, animacy, and size information increase sequentially — for instance, with colour information emerging from the first layer, animacy information from the third layer, and size information from the fifth layer — it suggests a trend of these layers becoming progressively deeper. Consequently, the emergence of colour, animacy, and size features was delayed. This could account for the progressively slower reaction times observed for colour, animacy, and size features in the behavioural results.

We extracted representational patterns for each picture in each layer, and then calculated Spearman’s correlations between the representational patterns of all pairs of pictures in each layer. This process yielded a total of eight representational similarity matrices (RSMs) (Figure 3A, only the lower triangles are shown). To discern feature-specific representations in each layer, the representational geometry in each layer was modelled as a weighted sum of models capturing the colour, animacy and size features (Figure 3B). The coefficients of the model RSMs represented the strength of feature-specific information. Model RSMs were constructed by assigning correlation distances of 1 to identical conditions and correlation distances of 0 to between-feature distances. Only the vectorized lower triangular regions of the RSMs (excluding the diagonal) were used. We used a permutation test to validate the statistical significance of the feature-specific representations (see Methods).

**Figure 3.**
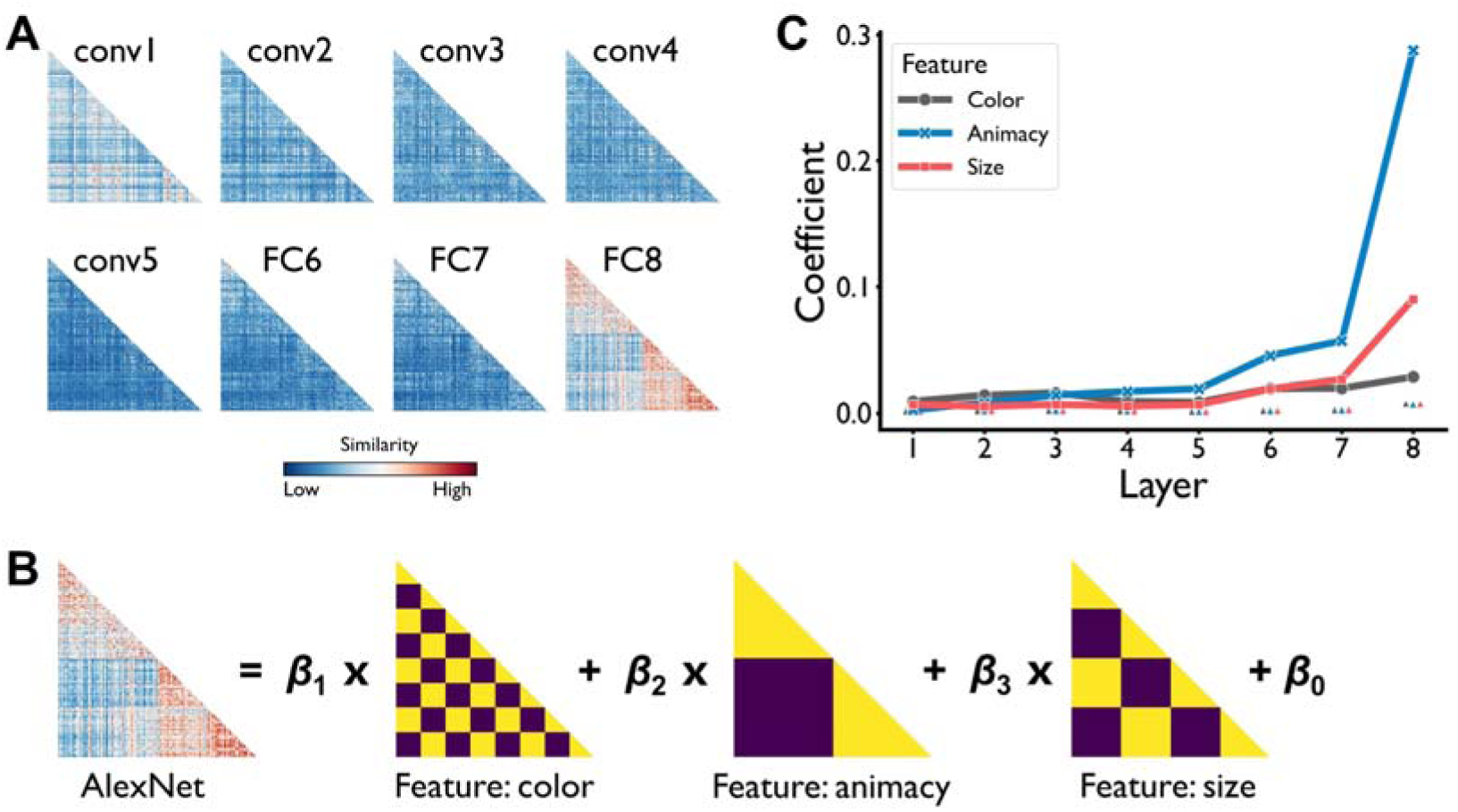
Representations of three features (colour, animacy and size) in the deep neural network AlexNet (stimulus set 1). **A. Representational similarity matrices.** The matrices were generated by correlating the activations of artificial neurons in each DNN layer. Only the lower triangular regions of the RSMs (excluding the diagonal) are shown. **B. Linear model.** The RSM of representational patterns of pictures in each layer was modelled as a weighted sum of models capturing the colour, animacy and size features. **C. Sensitivity of feature information.** This plot illustrates the sensitivity of feature information in each DNN layer. The small triangles, not connected by lines, represent the thresholds of feature-specific representations in each layer. Figure 3 was generated using data from stimulus set 1. The figure for the stimulus set 2 is available in the supplementary materials (Figure S2). conv: convolutional layer; FC: fully connected layer.

This test indicated that the coefficients for all three features consistently surpassed the corresponding thresholds obtained from the permutation test in each layer for both stimulus sets (Figure 3C, also see Table S1 & S2). These findings suggest that all eight layers contained information related to the three features. No hierarchical arrangement was observed where layers containing colour, animacy, and size features were positioned progressively higher. Therefore, we conclude that colour, animacy, and size features are processed in parallel within the deep neural network framework. Additionally, to determine the change rate of decision evidence across different layers of AlexNet, we calculated the average difference of coefficients between each pair of adjacent layers for each feature. The change rate of decision evidence for the perceptual feature colour is 0.003 for both stimulus sets, whereas the change rate of decision evidence for the conceptual features is 0.026 for stimulus set 1 and 0.027 for stimulus set 2 (averaged across the animacy and size features). These results indicate that the change rate of decision evidence is faster for conceptual features compared to perceptual features.

### The evidence accumulation process of three features

The AlexNet results indicated that the representation of these features began in the first layer and progressed to higher layers, and the increase in conceptual evidence across layers was faster than the increase in perceptual evidence. This seems contradictory to the reaction time (RT) data, where we observed a progressive increase in RTs for colour, animacy, and size features. To address this discrepancy, we further employed the drift-diffusion model (DDM) (Ratcliff, 1978; Ratcliff & McKoon, 2008a) to decompose the RT into different factors, including the rate of evidence accumulation, i.e., drift rate (δ), the non-decision time (τ) , and the boundary separation (α) that quantifies the amount of information or evidence required before a decision is made. Larger boundary separations, greater non-decision time and slower drift rates would lead to slower reaction times (Figure 4A).

**Figure 4.**
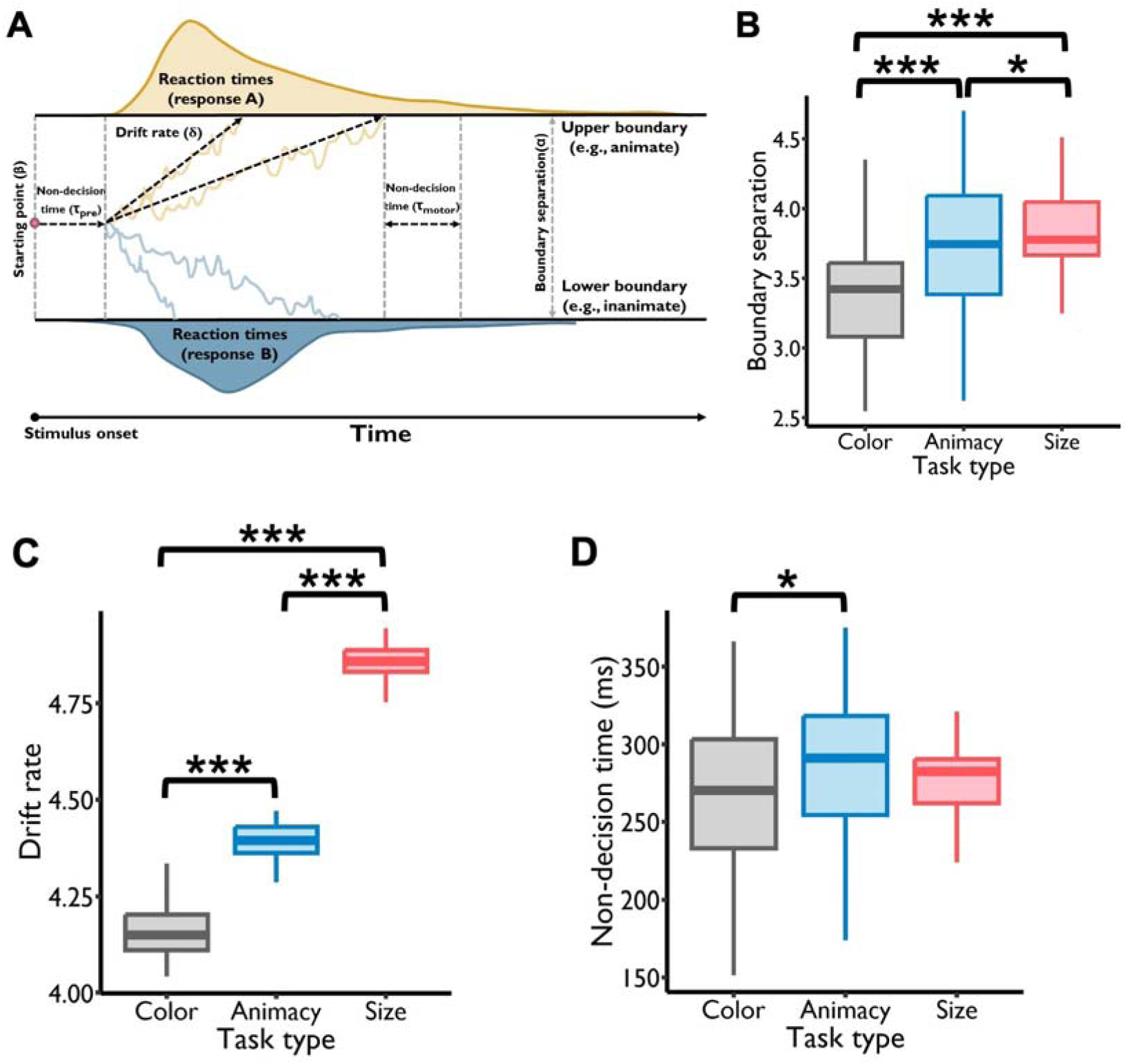
Results of the drift-diffusion model. **A. Graphical illustration of the drift-diffusion model.** At the onset of a trial, a participant’s information level commences at β. Over time, feature information accumulates until it reaches either the upper boundary (e.g., animate) or the lower boundary (e.g., inanimate), resulting in response A or B. The distance between the two decision boundaries is the boundary separation (α). The accumulation rate of feature information is represented by the drift rate δ. The time utilized for preprocessing before evidence accumulation and motor response after decision-making is the non-decision time τ. **B. Pattern of boundary separation.** A discernible pattern showcased increasing boundary separation along the color, animacy, and size tasks, and all of the comparisons were statistically significant. **C. Drift rate.** The drift rate (DR) of color, animacy and size features showed a significantly increasing pattern: DR_(color)_ < DR_(animacy)_ < DR_(size)_, and all of the comparisons were statistically significant. **D. Non-decision time.** The non-decision time of the color feature was significantly shorter than that of the animacy feature. *: *p* < 0.05, **: *p* < 0.01, ***: *p* < 0.001.

The mean boundary separation for the colour, animacy and size tasks was 3.39 (*SD* = 0.41), 3.71 (*SD* = 0.53) and 3.85 (*SD* = 0.26). Linear mixed-effects model revealed that task type significantly predicted the boundary separation (*F_2, 52_* = 46.47, *p* < 0.001). Post-hoc testing showed that the boundary separation for the size task was significantly larger than that for the colour (*t_52_* = 9.42, *p_corrected_* < 0.001) and animacy tasks (*t_52_* = 2.91, *p_corrected_* < 0.05), and the boundary separation for the animacy task was significantly larger than that for the colour task (*t_52_* = 6.50, *p_corrected_* < 0.001) (Figure 4B). In addition, the mean drift rate for the colour, animacy and size tasks was 4.16 (*SD* = 0.07), 4.39 (*SD* = 0.05) and 4.87 (*SD* = 0.05). Linear mixed-effects model revealed that task type significantly predicted the drift rate (*F_2, 52_* = 3115.56, *p* < 0.001). The drift rate (DR) of colour, animacy and size tasks showed a significantly increasing pattern: DR (colour) < DR (animacy) < DR (size), with all comparisons being statistically significant (animacy task vs color task: *t_52_* = 25.59, *p_corrected_* < 0.001; size task vs. color task: *t_52_* = 77.46, *p_corrected_* < 0.001; size task vs. animacy task: *t_52_* = 51.88, *p_corrected_* < 0.001) (Figure 4C). The mean non-decision time for the colour, animacy and size tasks was 265.47 ms (*SD* = 48.91 ms), 285.77 ms (*SD* = 52.06 ms) and 277.72 ms (*SD* = 23.83 ms). The results of the linear mixed-effects model showed that task type significantly predicted the non-decision time (*F_2, 52_* = 3.50, *p* < 0.05). Post-hoc testing showed that the non-decision time for the animacy task was significantly longer than that for the colour task (*t_52_* = 2.63, *p_corrected_* < 0.05). However, there were no significant differences in non-decision time between the colour and size tasks (*t_52_* = -1.59, *p_corrected_* = 0.26) or between the animacy and size tasks (*t_52_* = 1.04, *p_corrected_* = 0.55) (Figure 4D).

These results suggest that the differences in reaction times (RTs) among the three tasks were primarily due to the variations in boundary separation. Additionally, the non-decision time for the animacy feature was longer than that for the colour feature, which might also contribute to the longer RTs for the animacy task compared to the colour task. This confirms that larger boundary separations and longer non-decision times necessitate more time for decision-making. Interestingly, we found that the drift rate for the conceptual tasks was faster than for the perceptual tasks, aligning well with the AlexNet results. Nevertheless, this faster drift rate cannot completely compensate for the longer RTs caused by the larger boundary separations.

### Selective attention modulated the onset times of feature perception

Having revealed the parallel processing of perceptual and conceptual features, we further examined how selective attention could modulate these processes. To do this, we combined magnetoencephalography (MEG) and single-trial-based multivariate classification analyses to identify the onset and peak times of features in each trial of each task (color, animacy, and size). We hypothesized that target features would be represented earlier than non-target features during the perception stage.

Recognizing that attention may modulate different brain regions in distinct ways, all decoding analyses were carried out separately in four regions of interest (ROIs): the occipital, temporal, parietal, and frontal lobes. The sensors encompassed within each ROI were precisely defined using the MNE toolbox (Gramfort et al., 2013) (Figure S3). We used the learning stage as an independent localizer task to find the most informative time window (i.e., the peak bin) of each feature in each ROI (Liu et al., 2019; Wimmer et al., 2020). For each participant, four distinct classifiers were trained at each time point and ROI using data from the learning stage: one to classify the color category (colorful or gray), one for the animacy category (animate or inanimate), and two for the real-world size category (big or small) – one for animate objects and another for inanimate objects (Figure 5A). The decision to classify the real-world size category separately for animate and inanimate objects was informed by research indicating that real-world size drives differential responses only in the object domain, not the animate domain (Konkle & Caramazza, 2013). During the learning stage, we found significant clusters of color, animacy and size features in the occipital, temporal and parietal lobes (*P_corrected_* < 0.05). In the frontal lobe, we only found significant clusters of the animacy feature (*P_corrected_* < 0.05) (Figure S4). It should be noted that significant clusters for the size feature in animate objects were not identified during the learning stage. Consequently, the onset and peak times for the size feature during the perception and recall stages were exclusively calculated for inanimate objects. The peak latencies of significant clusters for the color, animacy, and size features during the learning stage were 305 ms, 515 ms, and 315 ms in the occipital lobe; 365 ms, 485 ms, and 445 ms in the temporal lobe; and 565 ms, 495 ms, and 445 ms in the parietal lobe. The peak latency of the animacy feature in the frontal lobe was 595 ms.

**Figure 5.**
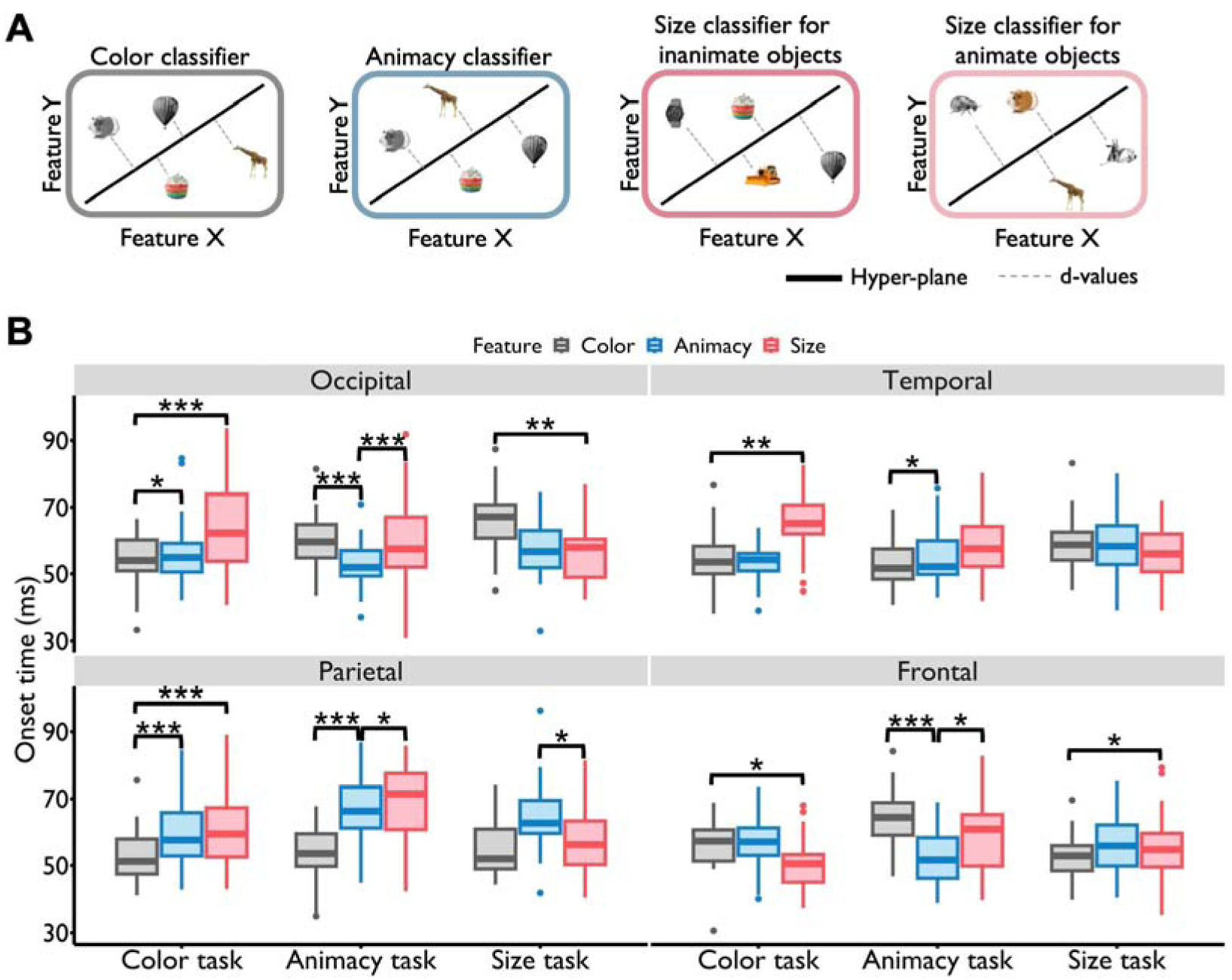
Modulation effects of selective attention during the perception stage. **A. Classifier training.** Four classifiers were trained in each ROI using data from the learning stage for each participant. These included one color classifier (colorful vs. gray), one animacy classifier (animate vs. inanimate), and two real-world size classifiers (small vs. big) for inanimate and animate objects. All classifiers were tested at each time point of each trial in the feature perception task. **B. Modulation Effects. 1) Comparing the animacy and size features:** In the occipital lobe, the animacy feature had significantly earlier onset times than the size feature during the animacy task, while no significant difference was observed in the size task. Notably, in the parietal lobe, the animacy feature had earlier onset times in the animacy task, whereas the size feature showed earlier onset times in the size task, reflecting a reversed pattern. **2) Comparing the color and animacy features:** In the occipital lobe, the color feature exhibited significantly earlier onset times than the animacy feature during the color task. Conversely, in the animacy task, the color feature had significantly later onset times compared to the animacy feature. **3) Comparing the color and size features:** In the occipital lobe, the color feature had significantly earlier onset times than the size feature in the color task, but significantly later onset times in the size task. Similarly, in the temporal lobe, the color feature showed earlier onset times in the color task, while no significant difference was observed between the two features in the size task. *: *p* < 0.05, **: *p* < 0.01, ***: *p* < 0.001.

To identify the most informative time window containing feature information, a 50 ms window was selected, which included the peak time point and four surrounding time points in each condition (i.e., peak bin). Utilizing these peak bins ensured that classifiers learned activation patterns of feature representations with the highest signal-to-noise ratio. Trained classifiers were then tested on data from each time point in each task type during the perception stage. The resulting time course of decoding performance for each feature in each trial, ROI, and participant was obtained.

Permutation and bootstrapping methods were employed to construct a null distribution for each time point. The 95^th^ or 5^th^ percentile of the null distribution served as the threshold for the corresponding time point (see Methods). A cluster with a duration of at least 20 ms, wherein all time points surpassed the corresponding thresholds, was considered to significantly contain feature information. The onset time of feature representation in a trial was defined as the first time point of the first significant cluster in that trial. This onset time represented the earliest time point at which feature information was detected in the brain. The peak time point, corresponding to when feature information was represented with the highest fidelity, was determined by selecting the time point with the highest d-value across all significant clusters in each trial. To investigate whether the onset or peak times of target features were earlier than those of non-target features during the perception stage, single-trial-based generalized linear mixed models (GLMM) were employed. The GLMM included task type (color, animacy, and size tasks), feature type (color, animacy, and size features), and their interactions as fixed effects. Participant ID, including the intercept, was modeled as a random factor.

For the onset times, we found that the interaction between task and feature was significant in the occipital (*F_4, 17104_* = 10.659, *p* < 0.001), temporal (*F_4, 16809_* = 6.663, *p* < 0.001), and parietal lobes (*F_4, 17066_* = 4.173, *p* = 0.002). Then we examined whether the onset times of (1) conceptual features (i.e., animacy vs. size) or, in a further step, (2) perceptual and conceptual features (i.e., color vs. animacy; color vs. size) were modulated by selective attention. We anticipated that the onset times of target features would precede those of non-target features. Additionally, if conceptual features were designated as target features, their onset times might precede those of perceptual features when the latter were non-target features.

For features belonging to the conceptual category, i.e., the animacy and size features, post-hoc analyses showed that in the occipital lobe, the onset times of the animacy feature (target feature) were significantly earlier than those of the size feature (non-target feature) in the animacy task (animacy feature vs. size feature: *t_17104_*= -4.019, *p_corrected_*< 0.001). However, in the size task, the onset times of these two features did not show a significant difference (animacy feature vs. size feature: *t_17104_* = -0.013, *p_corrected_* = 0.989) (Figure 5B). More importantly, in the parietal lobe, the onset times of the animacy feature were significantly earlier than those of the size feature in the animacy task (animacy feature vs. size feature: *t_17066_* = -1.969, *p_corrected_* = 0.049).

Conversely, a reversed pattern was found in the size task (animacy feature vs. size feature: *t_17066_* = 2.352, *p_corrected_* = 0.019) (Figure 5B). These results showed that selective attention made the onset times of the target feature occur earlier than those of the non-target feature.

For features belonging to different categories, i.e., the perceptual feature color and the conceptual feature animacy, post-hoc comparisons indicated that in the occipital lobe, the onset times of the color feature were significantly earlier than those of the animacy feature in the color task (color feature vs. animacy feature: *t_17104_* = -2.114, *p_corrected_* = 0.035). Conversely, in the animacy task, the onset times of the color feature were significantly later than those of the animacy feature (color feature vs. animacy feature: *t_17104_* = 4.901, *p_corrected_* < 0.001) (Figure 5B). These findings indicate that selective attention indeed influenced the onset times of both perceptual and conceptual features.

A similar pattern was observed for the color and size features. In the occipital lobe, the onset times of the color feature occurred significantly earlier than those of the size feature in the color task (color feature vs. size feature: *t_17104_* = -3.930, *p_corrected_* < 0.001), while in the size task, the onset times of the color feature occurred significantly later than those of the size features (color feature vs. size feature: *t_17104_* = 2.916, *p_corrected_* = 0.004) (Figure 5B). We also found that in the temporal lobe, the onset times of the color feature were significantly earlier than those of the size feature in the color task (color feature vs. size feature: *t_16809_* = -2.956, *p_corrected_* = 0.003), while in the size task, the onset times of the color and size feature did not differ (color feature vs. size feature: *t_16809_* = 1.138, *p_corrected_* = 0.255) (Figure 5B). These results highlight the important role of selective attention in modulating the onset times of features at different levels. Intriguingly, selective attention even led to earlier onset times for conceptual features compared to perceptual features.

### Phase coupling strength during task preparation correlates with earlier onset times of feature representations during perception

The above results showed that selective attention had a discernible impact on the onset times of both target and non-target features. Remarkably, this modulation effect was particularly pronounced in the occipital lobe, a sensory cortex. This observation raised the intriguing question of why the occipital lobe exhibited such a robust modulation effect. Previous research has posited that visual attention relies on brain networks comprised of higher-order regions, such as the temporal cortex attention network (Long & Kuhl, 2018; Ramezanpour & Fallah, 2022; Yeo et al., 2011). These networks are believed to support visual attention by influencing and assessing sensory representations within visual cortical areas (Desimone & Duncan, 1995; Egner & Hirsch, 2005; Gazzaley & Nobre, 2012; Serences & Yantis, 2006). Moreover, the increases in sensory responses (as estimated by spectral power) induced by selective attention were accompanied by synchrony between the higher- and lower-order regions (Baldauf & Desimone, 2014). Consequently, we assumed that the occipital lobe might not solely execute the entire modulation process on its own; instead, it might receive assistance from higher-order regions to achieve this modulation effect. To investigate whether the occipital lobe was influenced by other brain regions, we employed the phase slope index (PSI) to analyze phase coupling patterns between the occipital lobe and higher-order regions. We focused on four frequency bands: theta (4-8 Hz), alpha (9-13 Hz), beta (14-30 Hz), and gamma (31-100 Hz) during the task preparation phase, which occurred before each image was presented in the feature perception task. In this analysis, the occipital lobe served as the target region of interest (ROI), with the higher-order regions acting as seed ROIs. We then examined the relationship between the strength of this phase coupling and the onset times of feature representations in the occipital lobe. Our results revealed that the strength of phase coupling between the temporal and occipital lobes in the theta band significantly correlated with the onset times of target features, but not non-target features. Specifically, two significant clusters were identified in the theta band for each task type (*p_corrected_* < 0.001) (Figure 6A). In each task type, PSI values in the first significant cluster were positive, indicating a top-down flow of information from the temporal to the occipital lobe. Conversely, PSI values in the second significant cluster were negative, signifying a reversed information flow compared to the first cluster.

**Figure 6.**
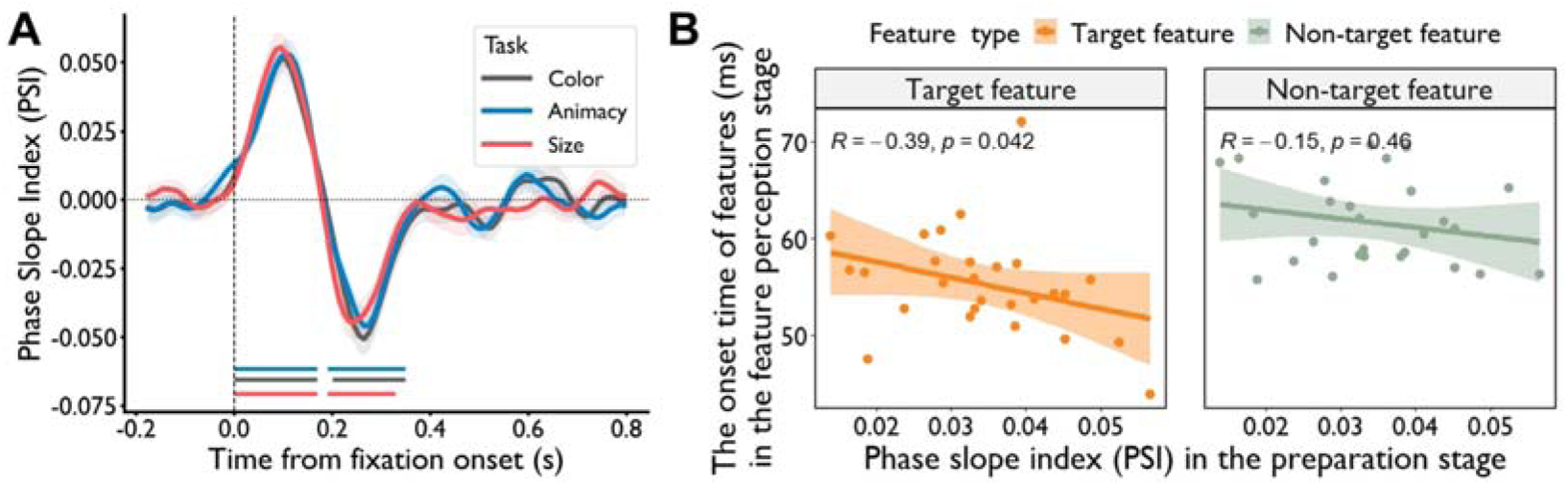
Phase coupling results in the feature perception task. **A. Phase coupling patterns during the task preparation stage.** During the task preparation stage, defined as the fixation period before the presentation of images, significant phase coupling in the theta band was observed between the temporal and occipital lobes for each type of task (*p_corrected_* < 0.001). Two distinct directions of coupling were identified: one from the temporal to the occipital lobe, and the other in the opposite direction. **B. Correlation with onset times.** A significant negative correlation was observed across subjects between the strength of temporal-occipital phase coupling in the theta band during the task preparation stage and the onset times of target features (*R* = -0.39, *p_corrected_* = 0.042), but not of non-target features (*R* = -0.15, *p_corrected_* = 0.46), in the occipital lobe during perception.

This suggested that, in the theta band, the information flow between the temporal and occipital cortex underwent a sequential top-down and bottom-up process during the task preparation stage. The phase coupling patterns between the temporal and occipital lobes across other frequency bands are illustrated in Figure S6.

Having obtained the phase coupling between the seed and target ROIs, we computed Pearson’s correlation between the phase coupling strength and the onset times of target and non-target features in the occipital lobe, respectively. This analysis aimed to explore whether a more robust phase coupling between the occipital lobe and higher-order regions during the preparation stage would be correlated with earlier onset times of target features rather than non-target features during the subsequent perception stage. Our findings revealed a significant negative correlation between the strength of phase coupling in the theta band between the temporal and occipital lobes and the onset times of the target features (*R* = -0.39, *p_corrected_* = 0.042; Figure 6B, left panel) while this was not the case for the non-target features (*R* = -0.15, *p_corrected_* = 0.46; Figure 6B, right panel) in the occipital lobe. This suggests that a stronger phase coupling between the temporal and occipital lobe in the theta band during the task preparation stage is associated with earlier onset times of target features in the occipital lobe during the subsequent perception stage.

### Selective attention modulates the peak times of feature retrieval

The previous analysis unveiled that selective attention modulates the onset times of features in the feature perception task, even leading to earlier onset times for conceptual features compared to perceptual features. In a subsequent analysis, we aimed to investigate whether selective attention could also influence the onset or peak times of feature representations during the memory retrieval stage. This inquiry is intriguing due to the distinct nature of visual perception and memory retrieval as two separate mental processes. Visual perception involves both outside-in and top-down processes, where individuals process visual inputs from the external world while also integrating expectations and predictions. In contrast, memory retrieval is primarily an inside-out process, involving the recall of information stored in the mind.

Additionally, past research has indicated that conceptual information is reconstructed faster than perceptual details during the memory retrieval stage, showcasing an inverted pattern compared to the object recognition process (Linde-Domingo et al., 2019; Mirjalili et al., 2021). The question of how selective attention’s modulation effects on feature representational timing vary across these two different mental processes remains unclear. Employing the same decoding and statistical analysis methods utilized in the feature perception stage, our findings revealed a modulation of selective attention on the peak times rather than the onset times of feature retrieval.

Specifically, results from the GLMM indicated that the interaction between task type (color, animacy, and size tasks) and feature type (color, animacy, and size features) was significant in the frontal lobe (*F_4, 8674_* = 5.697, *p* < 0.001), and at a trend level in the temporal lobe (*F_4, 8611_* = 2.189, *p* = 0.068).

For the conceptual features (animacy and size), post-hoc comparisons indicated that in the temporal lobe, the peak times of the animacy feature were significantly earlier than those of the size feature in the animacy task (animacy feature vs. size feature: *t_8611_* = -3.128, *p_corrected_* = 0.046), while the peak times of these two features showed no significant difference in the size task (animacy feature vs. size feature: *t_8611_* = -0.996, *p_corrected_* = 0.986) (Figure 7). For the perceptual feature color and conceptual feature animacy, post-hoc comparisons indicated that in the frontal lobe, the peak time of the color feature was significantly earlier than that of the animacy feature in the color task (color feature vs. animacy feature: *t_8674_* = -3.414, *p_corrected_* = 0.019), while this pattern was reversed in the animacy task (animacy feature vs. color feature: *t_8674_* = -3.121, *p_corrected_* = 0.047) (Figure 7). These results indicated that the perceptual feature (color) can be reconstructed with the highest fidelity earlier than the conceptual feature (animacy) in the frontal lobe. In the temporal lobe, the peak time of the animacy feature was significantly earlier than that of the color feature in the animacy task (animacy feature vs. color feature: *t_8611_* = -3.708, *p_corrected_* = 0.007), while in the color task, the peak times of these two features didn’t show a significant difference (animacy feature vs. color feature: *t_8611_* = -1.430, *p_corrected_* = 0.886) (Figure 7). For the color and size feature in the temporal cortex, the peak time of the size feature was significantly earlier than that of the color feature (size feature vs. color feature: *t_8611_* = -3.455, *p_corrected_* = 0.016), while there was no significant difference in the peak times between these two features in the color task (size feature vs. color feature: *t_8611_* = -1.885, *p_corrected_* = 0.624) (Figure 7).

**Figure 7.**
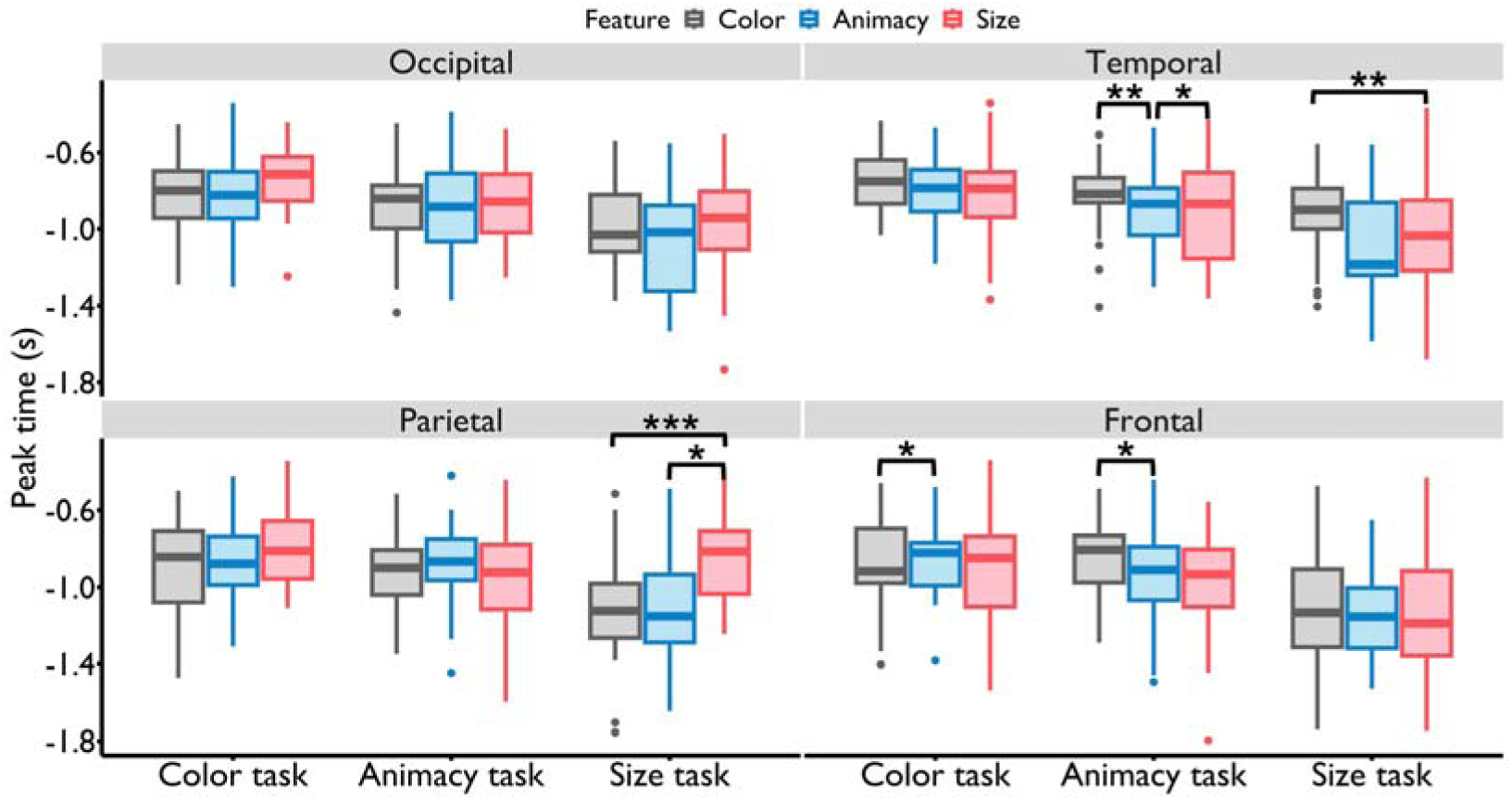
Modulation effects of selective attention on the peak times of feature representations in the feature retrieval task. **1) Comparing the animacy and size features:** In the animacy task, the peak time of the animacy feature in the temporal lobe occurred significantly earlier than that of the size feature. However, in the size task, the peak times of these two features showed no significant difference. **2) Comparing the color and animacy features:** In the frontal lobe, a reverse effect was observed where, in the color task, the peak time of the color feature occurred significantly earlier than that of the animacy feature, while in the animacy task, this pattern was reversed. In the temporal lobe, it was noted that in the animacy task, the peak time of the animacy feature was significantly earlier than that of the color feature. However, in the color task, the peak times of these two features showed no significant difference. **3) Comparing the color and size features:** In the size task, the peak time for the size feature in the temporal lobe was significantly earlier than for the color feature, while in the color task, the peak times for these two features exhibited no significant differences. *: *p* < 0.05, **: *p* < 0.01, ***: *p* < 0.001.

In contrast to the perception stage, our observations during the retrieval stage indicated that attention modulated the peak times rather than the onset times of feature representations. Notably, this modulation resulted in the target features being reconstructed with the highest fidelity earlier than the non-target features. The comparison between color and animacy features further suggested that attention modulations might facilitate a faster reconstruction of perceptual features compared to conceptual features.

Intriguingly, in contrast to the feature perception task where modulation effects primarily manifested in the occipital lobe, the feature retrieval task revealed modulation effects predominantly in the temporal and frontal lobes. We posited that information exchange between temporal and frontal lobes might commence as early as during the task preparation stage, specifically during the fixation period preceding the presentation of the spatial cue. Participants likely engaged in a preemptive object or target feature retrieval, facilitating rapid response upon the spatial cue’s appearance.

To scrutinize interactions between these lobes, we computed the phase slope index (PSI) across four frequency bands between the temporal (target ROI) and frontal (seed ROI) lobes, utilizing fixation period data before the spatial cue stage. Remarkably, significant clustering between these regions emerged in the beta band for each task type (color task: 225-305 ms, 365-515 ms; animacy task: 295-525 ms; size task: 415-535 ms) (*p_corrected_ <* 0.05) (Figure 8A). The values in each significant cluster were positive, signifying a prevailing direction of information flow from the frontal to the temporal lobe. A striking departure from this pattern became evident when scrutinizing the results of the perception preparation stage (Figure 8B). Notably, during this stage, no significant cluster featuring the same direction of information flow emerged as was seen in the retrieval preparation stage. Instead, we uncovered a significant cluster with information flow directed from the temporal to the frontal lobe, observed in the animacy (175-255 ms) and size task (115-195 ms) (*p_corrected_ <* 0.05).

**Figure 8.**
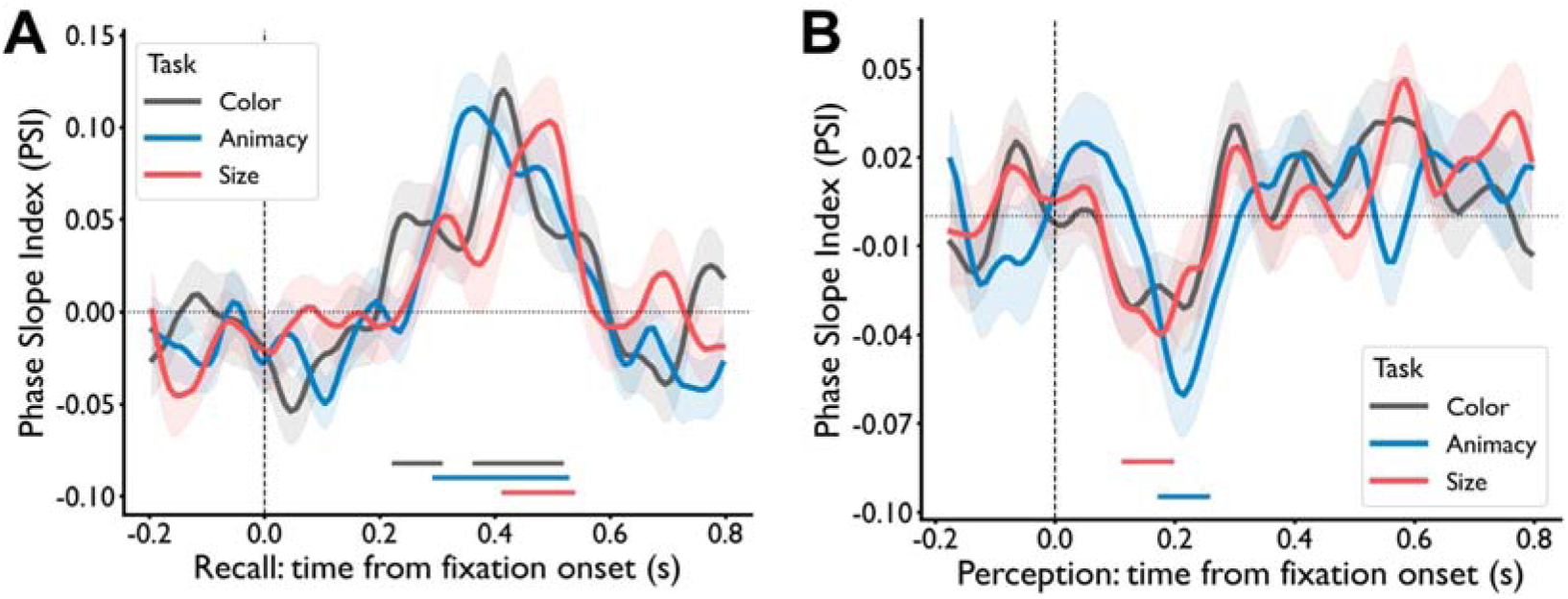
Phase coupling between temporal and frontal lobes in the beta band during the preparation stage. **A.** In the feature recall task, a significant cluster of phase coupling from the frontal to the temporal lobe was observed in each task type (color, animacy and size task) (*p_corrected_* < 0.05). **B.** In the feature perception task, a significant cluster of phase coupling from the temporal to the frontal lobe was found in the animacy and size tasks (*p_corrected_* < 0.05).

These nuanced results underscore the specificity of the frontal-to-temporal information flow during the task preparation stage of the retrieval task rather than the perception task.

## Discussion

The question of whether the flow of information across different feature levels within objects adheres to a fixed hierarchical order during visual perception and retrieval stages is intriguing. Our findings suggest that the representation times for different levels of features, such as perceptual and conceptual features, can indeed be modulated by selective attention. Specifically, during the visual perception phase, conceptual features can be detected earlier than perceptual features. Conversely, during the memory retrieval phase, perceptual features can be reconstructed with the highest fidelity before conceptual features. This study offers valuable insights into the temporal modulation effects of selective attention, significantly advancing our understanding of the nature of processing hierarchy and how attentional processes differentially influence the timing of feature detection and reconstruction in visual perception and memory retrieval, respectively.

Our study, utilizing the deep neural network AlexNet (Krizhevsky et al., 2017), demonstrated the parallel processing of color, animacy, and real-world size features, validating the sequential order-free nature of their processing. Furthermore, employing the drift-diffusion model (Ratcliff, 1978; Ratcliff & McKoon, 2008b), we observed a sequential increase in boundary separation for color, animacy, and size features, corresponding with their reaction times. Our findings align with previous theoretical and empirical research. Theoretical models have suggested the simultaneous processing of object information across different categorization levels, with varied evidence accumulation time courses (Kravitz et al., 2013; Mack & Palmeri, 2011).

Empirical research has shown early onset latencies for decoding accuracy across various categorization levels (48-70 ms), with higher-level features peaking later than lower-level features (Carlson et al., 2013; Cichy et al., 2014).

To further explore whether the temporal dynamics of perceptual and conceptual feature perception can be modulated by selective attention, we employed single-trial-based decoding analysis, a highly sensitive method for detecting representational times (Linde-Domingo et al., 2019). As emphasized by Linde-Domingo et al., 2019, many factors can obscure differences in timing between conditions when averaging decoding traces across trials and subjects. For instance, there was substantial variance in reaction times among participants. Similarly, there were significant variances across trials within the same subject, as factors such as retrieval speed, mental state, and other conditions fluctuate across trials. Averaging decoding traces across trials and subjects would reduce the effective feature information, making it difficult to detect timing differences between conditions. Therefore, a single-trial-based method was considered to be more sensitive in capturing such differences. Based on this method, our results revealed that in the visual perceptual task, directing attention to perceptual features (e.g., color) resulted in their onset times preceding those of conceptual features. Interestingly, when attention was shifted to conceptual features (e.g., animacy and real-world size), the onset times of these conceptual features significantly preceded those of perceptual features (see Figure 5). These findings suggest that selective attention enables the earlier detection of target features over non-target features. Furthermore, in conceptual tasks, attention can orchestrate a scenario where conceptual features took precedence over perceptual features.

Numerous studies have demonstrated the modulation effects of selective attention on various aspects of information processing and brain responses, including activation strength (Baldauf & Desimone, 2014; Maunsell & Treue, 2006), representational fidelity (Grootswagers et al., 2021; Long & Kuhl, 2018) , dimensions (Sheng et al., 2022) and distance (Nastase et al., 2017). Our findings on the adjustable onset times of perceptual and conceptual feature perception underscore the dynamic and flexible nature of visual perception from a new perspective. This challenges the notion of a fixed sequence where perceptual features invariably precede conceptual features (Clarke et al., 2013; Linde-Domingo et al., 2019; Mirjalili et al., 2021). These results also contribute to the intense debates on the nature of perception, which center on passive (purely bottom-up) versus active accounts (Firestone & Scholl, 2016; Treisman, 1980). Notably, theories of active sensing, dating back to Gibson’s seminal work in the 1950s, propose that perception is not merely a passive reception of sensory inputs but is fundamentally driven by internal goals and motivations (Gibson, 1950; O’Regan & Noë, 2001; Summerfield & De Lange, 2014). According to these active accounts, task demands and attention play crucial roles in shaping how information is processed and represented in the brain (Desimone & Duncan, 1995; Engel et al., 2001; Nobre & Kastner, 2014). This perspective suggests that our perceptual experiences are not solely dictated by external stimuli but are also significantly influenced by our cognitive states and objectives.

It is noteworthy that the modulation effect of selective attention was observed in the occipital lobe for all three feature pairs during the visual perception stage. Previous research has shown that top-down factors can influence stimulus representations in early visual areas (Baldauf & Desimone, 2014; Ester et al., 2016; Jehee et al., 2011; Lu et al., 2015; Sheng et al., 2022; Sprague & Serences, 2013; Xue et al., 2013; Zheng et al., 2018). Moreover, cognitive intentions, such as expectations and mental readiness embedded in top-down processes, may precede the presentation of a stimulus. This intentional processing guides bottom-up sensory processing by reallocating attention based on subjective expectations, thereby enhancing the efficiency of perceptual identification (Baldauf & Desimone, 2014; Córdova et al., 2016; Sakai, 2008; Siegel et al., 2008). Our findings replicate and extend these studies by demonstrating that selective attention can modulate the speed of feature accumulation, thereby influencing the temporal dynamics of visual perception.

To understand how this modulation was achieved, we explored the phase coupling between the occipital lobe and higher-order brain regions during the task preparation stage, when stimuli were not presented. In the theta band, we observed an initial information flow from the temporal to occipital lobe during the first 200 ms, followed by a backward information flow in the subsequent 200 ms. Further analysis revealed that the strength of phase coupling in this initial top-down and subsequent bottom-up cluster during the task preparation stage correlated with the onset times of target features, rather than non-target features, in the occipital lobe during the perception stage. Specifically, stronger phase coupling between the occipital and temporal lobes in the theta band was associated with earlier onset times of target features but not non-target features. This finding emphasizes the role of inter-regional interactions during the preparation stage in promoting the speed of evidence accumulation for target features in visual perception tasks.

In addition to perception, our results demonstrated that selective attention could modulate evidence accumulation during retrieval so that conceptual features do not always peak earlier than perceptual features (Linde-Domingo et al., 2019; Mirjalili et al., 2021). We observed a reversal of the peak times for perceptual and conceptual feature retrieval in the frontal lobe, highlighting the significant role of frontal regions in selective attention. Consistent with our findings, previous research has emphasized the crucial role of frontal regions in prioritizing items in visual working memory, selecting and integrating task-relevant features, and maintaining working memory flexibility while avoiding interference (Buschman & Kastner, 2015; Chatham et al., 2014; Nee & Jonides, 2008; A. C. Nobre et al., 2004). These regions are essential for the selection and integration of task-relevant features (Mante et al., 2013) and for maintaining the flexibility of working memory, thereby preventing interference from competing representations (Bouchacourt & Buschman, 2019).

In addition to the frontal lobe, we found that the temporal lobe also exhibits a selective attention modulation effect, suggesting that this effect extends to lower-level regions in the processing hierarchy. Previous research has shown that the frontal cortex, especially the prefrontal cortex, has reciprocal anatomical connections with the inferior temporal cortex (ITC) (Gerbella et al., 2010; Saleem et al., 2014). These connections exert top-down signals that modulate the visual neural responses of the ITC (Fuster et al., 1985) and induce activity related to memory retrieval (Tomita et al., 1999; Zhou et al., 2023). Our phase coupling analysis revealed a significant positive cluster in the beta band between the frontal (seed ROI) and temporal (target ROI) lobes, indicating information flow from the frontal to the temporal lobe. Notably, this directional information flow was absent during the preparation stage in the feature perception task, indicating task-specific information flow. These findings illustrate that the brain engages in distinct preparatory processes depending on the task type, whether it is a feature perception or retrieval task. These differences manifest in various ways, including the regions involved in phase coupling and the frequency bands through which information is communicated.

In addition to the differences in modulated brain regions and preparatory states across the perception and memory stages, it is particularly intriguing that the timing metrics influenced by selective attention also vary between these two stages. Our study identified an interesting discovery, showing that selective attention modulates the onset time of feature representations during the perception stage, whereas it modulates the peak time of feature representations during memory retrieval. Regarding why selective attention modulated the onset time of feature perception but only the peak time of feature retrieval, prior research provides valuable insights. In the process of perception, subjects do not need to identify the specific object to categorize certain features, such as animacy or size (Wang et al., 2022). However, during memory retrieval in our study, when participants were tasked with recalling a specific feature of an object at a cued location, they first needed to retrieve the object itself from that location. Previous research has shown that when an object is recalled, its binding features are often retrieved spontaneously, even when participants are not explicitly instructed to recall those features (Bone et al., 2020; Linde-Domingo et al., 2019).

Because these features are retrieved automatically during recalling an object, this may make selective attention less capable of modulating the onset time of their reactivation, meaning it may not determine when these features are first represented during retrieval. In contrast, during feature perception, selective attention can modulate onset times because individual features can be represented before the object is recognized, providing the possibility for attention to influence when these features are first represented. However, selective attention did make a difference. According to our results, it ensured that the target feature was represented with the highest fidelity before non-target features. This might suggest that selective attention accelerated the evidence accumulation speed for target features after both target and non-target features were spontaneously reactivated. In summary, the reason selective attention modulated different time metrics during the feature perception and feature retrieval stages may lie in the fact that the former did not require the representation of the object itself (Wang et al., 2022), allowing for the possibility of selective modulation of the onset time of feature perception. In contrast, the latter required the representation of the object itself in our experiment, leading to the spontaneous reactivation of its associated features (Bone et al., 2020; Linde-Domingo et al., 2019), which prevented selective attention from modulating the onset time of feature retrieval. This distinction warrants further investigation in future research to better understand the underlying mechanisms of selective attention across different cognitive stages.

Several questions require further examination. First, although MEG provides excellent temporal resolution, its spatial resolution is limited compared to other neuroimaging techniques like fMRI. Future research could conduct parallel MEG and fMRI studies to gain more detailed insights into both the temporal and spatial dynamics of the modulation effect of selective attention on feature processing. Second, while phase coupling analysis offers valuable insights into neural connectivity, interpreting these interactions remains complex. Employing additional methods, such as direct neural manipulation (e.g., Transcranial Magnetic Stimulation), in future research could better elucidate the causal relationships between neural connectivity and information processing. Finally, although computational models have been applied to understand other effects of selective attention, future studies could utilize computational models to further understand the role of selective attention in modulating the processing hierarchy.

To conclude, this study suggests that perceptual and conceptual features are processed in parallel, and selective attention modulates the temporal dynamics of their processing during both perception and memory retrieval tasks. This modulatory effect is correlated with the phase coupling between the temporal and occipital lobes in the theta band during perception and between the frontal and temporal lobes in the beta band during retrieval. These results underscore the dynamic and flexible nature of attentional modulation in cognitive processing.

## Methods

### Participants

Thirty healthy volunteers with normal or corrected-to-normal visual acuity and no history of psychiatric or neurological disorders participated in the study. Three of the participants were too drowsy to complete the experiment, so the analysis was conducted on the remaining 27 participants (8 female; mean age = 21.70±2.37 years).

All experiments were carried out in accordance with the Declaration of Helsinki. All participants provided written informed consent prior to the experiment, which was approved by the Research Ethics Committee at Peking University and the State Key Laboratory of Cognitive Neuroscience and Learning at Beijing Normal University in China.

### Stimuli

To scrutinize the processing hierarchy during both the perception and retrieval stages, it was crucial to independently manipulate the perceptual and conceptual features of the presented objects. In our experiment, objects varied along three orthogonal dimensions: one perceptual dimension (color), where objects were presented either in a colorful or grayscale format, and two conceptual dimensions — animacy and real-world size. Objects were categorized as either animate or inanimate (animacy dimension) and as either big or small (real-world size dimension) (Figure 1C).

We utilized 192 distinct images of everyday objects and common animals in the primary experiment, with an additional 24 images used in the practice session. Three types of features were introduced: color, animacy, and real-world size. For the color feature, half of the pictures were presented in grayscale, and the remaining half in color. To control for potential color-related biases, two sets of pictures were created with reversed color assignments between the sets (e.g., a colorful cupcake in stimulus set 1 and a gray cupcake in stimulus set 2) (Figure 1C). Half of the participants were exposed to stimulus set 1, while the other half encountered stimulus set 2.

The two animacy levels (animate and inanimate) and size levels (big and small) were evenly distributed within each level of the color feature (colorful and gray). The criteria for determining real-world size were based on a reference box measuring approximately 40 cm (length) × 18 cm (width) × 17 cm (height) in the experiment room. Objects that fit into this box were classified as small, typically handheld items, while those that didn’t fit were deemed big and could support a human (e.g., chair-sized or larger).

The 192 objects were divided into three groups (group 1, group 2, and group 3) for the three tasks (color, animacy, and size categorization tasks). Six assignment plans were implemented to ensure all combinations of stimulus groups and tasks were covered. In each stimulus group, an equal number of images were allocated to each level of every feature. For instance, the number of images representing animate and inanimate objects was the same. All images were presented at the center of the screen, with a rescaled size of 198×198 pixels.

### Procedure

The experiment comprised two parts: the feature retrieval task and the feature perception task. Before each task, participants received verbal instructions and practiced to become familiar with the experimental procedure. The feature retrieval task consisted of eight blocks, each containing three mini-blocks (24 mini-blocks in total). Each mini-block consisted of a learning stage and a recall stage. During the learning stage, a jittered fixation cross (1-1.5 s from a uniform distribution) was followed by a series of objects presented in pseudorandom order within the eight cells surrounding the central cell (no object was shown in the center cell). Each object was displayed for three seconds, with an inter-stimulus interval jittered around 1-1.5 seconds from a uniform distribution. Participants were instructed to learn the object-location associations in preparation for a memory test, with no responses required during the learning stage. The learning process in each mini-block was repeated three times consecutively. Objects of different types occurred with equal probability across all cells during the entire feature retrieval task.

Following the learning stage, participants engaged in the feature retrieval and object recognition tasks, where they were prompted to recall the features of the cued objects and distinguish target items from lures (Figure 1A). At the beginning of the retrieval stage, a task cue (indicating the color, animacy, or size task) was displayed for four seconds, instructing participants on which feature of the cued object they needed to retrieve (the target feature). A centrally displayed fixation cross appeared for 1 to 1.5 seconds, serving as the task preparation stage, before the beginning of each retrieval trial. Subsequently, a white square, identical in size to the pictures presented during the learning stage, appeared in any one of the eight cells. Participants were required to retrieve the target feature of the cued object and press the button to provide their answers upon successful recall. For example, in the animacy task, participants had to categorize whether the object in the corresponding location was animate or inanimate. Following this, three pictures were presented on the screen: a target picture shown during the learning stage and two lure pictures resembling the target picture.

Participants were tasked with choosing the target picture by pressing the corresponding button. Lure pictures ensured that participants remembered the visual details of the pictures and that the mental process of feature retrieval was based on the retrieved images. Each feature retrieval and object recognition trial allowed participants up to ten seconds to respond. Rest breaks were provided after each block, and the feature retrieval task lasted approximately one hour.

Following the feature retrieval task, participants took a break before commencing the feature perception task, which also utilized a block design (Figure 1B). Each block started with a task cue displayed for 4 seconds, directing participants on which feature to categorize (the target feature). During each trial, a jittered fixation cross was displayed centrally for 1 to 1.5 seconds during the task preparation stage. After this, a random picture appeared in one of the eight cells. Participants were required to rapidly and accurately categorize the object based on the target feature, pressing a button to provide their answers within 1.5 seconds of seeing the picture. Once the key response was made, the picture vanished from the screen, leaving a blank nine-cell grid. If the key response was not submitted within 1.5 seconds of the picture onset, the picture would also disappear. The combined duration of the picture and the blank nine-cell grid was 2 seconds. Objects of different types occurred with equal probability in each cell across the entire feature perception task. The entire feature perception task lasted approximately half an hour.

Throughout both the feature retrieval and perception tasks, key responses were counterbalanced across participants. For instance, one participant might use the left key for the colorful level, while another might use the right key for the same purpose. However, within each participant, the key response assignments remained consistent throughout the experiment. This approach aimed to mitigate the potential switching cost of key responses, which could otherwise prolong reaction times. Our objective was to obtain accurate reaction times for the feature categorization task and minimize the influence of task-switching costs. All stimuli were presented using PsychoPy (Peirce et al., 2019).

### MEG data acquisition

MEG data were acquired using a 306-sensor (204 planar gradiometers, 102 magnetometers) Elekta Neuromag MEG system (Helsinki, Finland) at Peking University, Beijing, China. Participants completed the MEG experiments inside a sound-attenuated, dimly lit, and magnetically shielded room. The head position inside the MEG helmet was continuously monitored using four head position indicator (HPI) coils. A 3D digitizer was used to record the location of the HPI coils and the general head shape relative to three anatomical fiducials (nasion, left and right preauricular points). Both horizontal and vertical electrooculograms (EOGs) were recorded. MEG data were recorded at a sampling frequency of 1000 Hz and bandpass filtered between 0.1 and 330 Hz.

### Deep neural network representations

To investigate whether the three features were processed in parallel or if some features were processed ahead of others, we used the deep neural network "AlexNet" (Krizhevsky et al., 2017). AlexNet consists of eight layers—five convolutional layers and three fully connected layers—which simulate the hierarchical structure of neurons along the ventral visual stream. Generally, the early layers of AlexNet process early visual features, such as colors, contrasts, and frequencies, while the deeper layers process higher-order visual representations, such as the surface structure of objects or body parts of animals. The AlexNet used in this study was pretrained using the ImageNet dataset (Deng et al., 2009).

We extracted the features of each picture from each layer of AlexNet and calculated the similarity between the features of every two pictures, excluding the similarity between the same picture. All correlation values were Fisher Z-transformed before further analyses. To obtain the feature-specific representations, we modeled the feature representational geometry of pictures in each layer of AlexNet as a weighted sum of models capturing the color, animacy and size features (Figure 3B). A category similarity matrix was constructed by assigning ones (maximum similarity) to elements of the matrix that corresponded to pairs of objects belonging to the same category (e.g., both were animate) and zeroes (minimum similarity) for pairs of objects from different categories (e.g., one is animate and the other is inanimate). After constructing the representational pattern similarity matrix (RSM) for each layer and three category similarity matrices, we performed regression-based representational similarity analyses to determine the contribution of category similarity to the representational patterns of AlexNet. Only the vectorized lower triangular regions of the RSMs (excluding the diagonal) were used for further analyses. The representational similarity in each layer was modeled as a linear combination of category similarity for the color, animacy, and size features (Figure 3B). This produced beta weights for the color, animacy and size features, respectively, for each layer.

We also used the permutation test to examine the statistical significance of the feature-specific representations (i.e., the beta values). For each feature in each layer, a null distribution was created through 1000 permutations. The threshold for feature-specific representations was determined by selecting the 95th percentile from this null distribution. During each permutation, the correspondence between the picture labels and representational patterns in each layer was shuffled. Subsequently, the feature-specific representations were computed using the same procedure as for the actual labels.

### The drift-diffusion model

To explore the evidence accumulation process of three features (color, animacy and real-world size) during the perception stage, we used the drift-diffusion model to extract the parameters characterizing the evidence accumulation process of features from the reaction times. Drift-diffusion models are widely used for predicting participants’ choices and reaction times (RT) during two-choice decision-making tasks (Ratcliff, 1978; Ratcliff & McKoon, 2008b; Stone, 1960). In the drift-diffusion model, it is assumed that participants accumulate evidence for one choice over another. The rate of evidence accumulation within a trial is called the drift rate (δ) (Figure 4A). Evidence is accumulated toward either the upper boundary (e.g., animate) or the lower boundary (e.g., inanimate), corresponding to two respective responses.

The boundary separation (α) represents the distance between the two decision boundaries, indicating the amount of evidence needed to make a decision. The starting point (β) reflects the initial position of evidence accumulation and indicates response bias toward one of the two response boundaries. The non-decision time (τ) is the portion of the reaction time not dedicated to the decision-making process, typically including preprocessing time and motor response time. The drift-diffusion model was implemented using the *brms* package in R (Bürkner, 2017). Only correct trials were used to fit the model. After fitting the model, we extracted the boundary separation, drift rate and non-decision time for each task and participant.

### MEG pre-processing

Raw MEG signals were cleaned using the signal space separation (Taulu & Simola, 2006) method provided by MaxFilter to suppress magnetic interferences and interpolate bad MEG sensors. MEG data were pre-processed using the MNE toolbox (version 1.1) (Gramfort et al., 2013). To remove line noise the data were band-stop filtered at 50 Hz and its harmonics. Muscle artifacts were automatically detected by the MNE function *annotate_muscle_zscore* with default parameters, and the annotations of muscle artifacts were then visually inspected by the experimenters. Signals annotated as muscle artifacts were excluded from further analysis.

Data recorded during the learning stage were epoched between -500 ms and +3000 ms relative to the picture onset. During the perception stage, data of feature categorization were epoched between -500 ms and +1000 ms relative to the picture onset. The epochs were baseline-corrected based on the pre-stimulus signal (-500 ms to onset). During the retrieval stage, data for feature retrieval were epoched between -4000 ms and +1000 ms relative to the key response. Since the post-response signal during retrieval might still contain task-relevant (i.e., feature-specific) information, we baseline-corrected the signal based on the whole trial. The data during the fixation cross before the feature perception (or retrieval) were epoched between -200 ms and 800 ms relative to the fixation onset. These epochs were baseline-corrected based on the pre-fixation signal (-200 ms to onset). Additionally, epochs with excessive noise were discarded using the autoreject toolbox (Jas et al., 2017). Independent component analysis was then used to remove eye-blink, eye movement, and electrocardiogram (ECG) artifacts. To increase the signal-to-noise ratio (SNR) for multivariate decoding, the pre-processed MEG time courses were smoothed using a Gaussian kernel with a full width at a half-maximum of 10 ms and down-sampled to 200 Hz.

### Time-resolved multivariate decoding

Before conducting the decoding analysis, we used a smooth window with a 50 ms length and a 10 ms step, averaging signals in each window to further increase the SNR. Our primary focus was on investigating the selective attention modulation effect on feature representations during the perception and retrieval stages. The learning task functioned as a functional localizer, enabling the training of independent feature classifiers without confounding effects related to motion. For each participant, four distinct classifiers were trained at each time point and region of interest (ROI) using data from the learning stage. These classifiers included one for color category classification (colorful vs. gray), one for animacy category classification (animate vs. inanimate), and two for real-world size category classification (big vs. small)—one for animate objects and another for inanimate objects (Figure 5A). The reason for classifying the real-world size separately for animate and inanimate objects was based on research indicating that real-world size drives differential responses only in the object domain, not the animate domain (Konkle & Caramazza, 2013). Therefore, we assumed that the representational patterns of real-world size for animate and inanimate objects might differ. The feature perception epochs of the same object were averaged. For both the epochs of feature perception and retrieval, the pre-processed amplitudes of channels in each ROI at each time point were whitened and then used as features for classifiers. The whitening process involved using a standard scaler that z-scored each channel at each time point across trials. Classification using logistic regression models was then performed separately for each time point, ROI, and participant, with a stratified 8-fold cross-validation approach. The logistic regression model parameters were set to their default values as provided by the Scikit-learn package (King et al., 2016), using L2 regularization, a tolerance for stopping criteria as 1e-4, an inverse of regularization strength (C) as 1, and the ‘lbfgs’ solver (Limited-memory Broyden–Fletcher–Goldfarb–Shanno) for optimization. The decoding performance during the learning stage was indicated by the Area Under the Receiver Operating Characteristic Curve (ROC_AUC). We used cluster-based permutation analyses (Maris & Oostenveld, 2007), which intrinsically correct for multiple comparisons, to test the significance of feature representations during the learning stage in each ROI.

After identifying the significant cluster of each feature in each ROI at the group level (averaged across participants) during the learning stage, the time point of peak ROC_AUC within the significant cluster and its surrounding four time points (e.g., if the peak time point was the 78^th^, the surrounding time points were the 76^th^, 77^th^, 79^th^, and 80^th^) were considered the most informative time window (i.e., peak bin) carrying the strongest feature information. The averaged and standardized amplitudes in each peak bin were used to train the feature classifiers, which were then tested at each time point in each task during the perception and retrieval stages to detect feature information without any motor confounding effects. For example, the animacy classifier trained during the learning stage in each ROI was tested in the three tasks (the color, animacy and size tasks) respectively during the perception and retrieval stages in the corresponding ROI. Cross-validation was not used for across-stage analyses as no inferences were made based on the decoding performance in the learning stage itself.

To detect the onset and peak times of feature representations during the perception stage, following the approach of Linde-Domingo et al., 2019, we conducted classification analyses at a single trial level. As emphasized by Linde-Domingo et al., 2019, many factors could obscure differences in timing between conditions when averaging latencies across trials, such as variance in processing speed. Therefore, the single-trial-based method was considered more sensitive to differences in timing between conditions. We tested the trained classifiers at each time point of each trial, resulting in a total of 16 decoding performance time courses for each participant and each trial: 4 ROIs (occipital, temporal, parietal, and frontal cortex) by 4 features (color for all objects, animacy for all objects, and size for inanimate and animate objects respectively). Unlike Linde-Domingo et al., 2019, we used both the peak latency of significant clusters of each trial as the time index to measure the time lag between different features and the onset (i.e., the first time point) of the first significant cluster of each trial. We posited that the onset time signifies the initiation of feature representations, capturing the very inception of when these features begin to manifest in the brain. Detecting this initial time point is crucial for understanding when these features first come into representation. On the other hand, the peak latency signifies the time at which a feature is represented with the highest fidelity. In this context, the onset time allows us to discern whether tasks induce the earlier detection of target features compared to non-target features in the brain. Meanwhile, the peak time helps us understand whether target features are represented with highest fidelity before non-target features. This dual approach provides a comprehensive understanding of the temporal dynamics of feature representations modulated by selective attention.

The decision value (i.e., d-value) is an appropriate index for characterizing decoding performance at the single-trial level (Linde-Domingo et al., 2019; Ritchie et al., 2015b). The d-value has both negative and positive signs, indicating the feature level to which the observation was classified (e.g., - for animate and + for inanimate). The absolute d-value at each time point represents the distance to the hyper-plane that divides the two feature levels (e.g., animate and inanimate), with the hyper-plane being 0 (Figure 5A). This distance indicates the classifier’s confidence in assigning a given object to a specific feature level at that time point. For extracting significant clusters carrying feature information for each single trial, the cluster-based permutation analysis based on multiple trials is not applicable. Therefore, we compared d-values at each time point in each trial with the threshold of the corresponding time point. A time window of at least T ms, where all the time points surpassed the corresponding thresholds, was considered a significant cluster. Details about null distributions are provided in the next section, “Generating empirical null distributions for classifiers.”

### Generating empirical null distributions for classifiers

To detect the significant clusters of feature representations at the single-trial level, we followed the method in Linde-Domingo et al., 2019, using permutation and bootstrapping to generate null distributions for each time point of each feature level in each task type. We then chose the 95^th^ or 5^th^ percentile of each null distribution as the threshold for the corresponding time point. In each trial, a time window of at least T ms where all the time points surpassed the corresponding thresholds was considered a significant cluster. The first time point of the first significant cluster was considered the earliest time point when the feature was represented in this trial, i.e., the onset time. The time point with the highest d-value across all significant clusters in each trial was considered the time point when the feature was represented with the highest fidelity, i.e., the peak time.

For the permutation analyses, we randomly shuffled the feature level labels (e.g., shuffled the colorful and gray labels) of trials during the learning stage at each permutation and carried out the same training procedure as for the real data. The classifiers were then tested on the data of three tasks during the perception and retrieval stages using the same test procedures as for the real data. This procedure was conducted 150 times independently per participant. For each participant, ROI, task, and feature type, there was a total of 151 classification outputs for each trial of the perception and retrieval stages respectively: one using the real labels and the remaining using the randomly shuffled labels when training the classifiers. The real data were included to make our subsequent bootstrapping analyses more conservative, since under the null hypothesis, the real classifier output could have been obtained just by chance.

Then the bootstrapping approach was used to estimate the classification chance distribution. For each ROI, task, and feature type, we randomly selected one of the 151 classification outputs per participant in each bootstrapped repetition. Trials in this classification output were divided into two groups based on their feature levels, and the d-value time courses of each feature level were averaged across trials. As a result, for each ROI, task, and feature type in each bootstrapped repetition, each participant had two d-value time courses—one feature level each (e.g., one for the animate level and the other for the inanimate level). The d-value time courses for each feature level were then averaged across participants. This procedure was repeated with replacement 10000 times. Therefore, for each feature level, each time point had a null distribution consisting of 10000 values. For levels with positive d-values (i.e., gray, inanimate, and small), we selected the 95^th^ percentile of each null distribution as the threshold for the corresponding time point. For levels with negative d-values (i.e., colorful, animate, and big), we selected the 5^th^ percentile of each null distribution as the threshold for the corresponding time point. This decision was made because the absolute value of the 5^th^ percentile would be higher than that of the 95^th^ percentile for negative values, indicating stronger classification confidence. In each trial, a time window of at least T ms where all the time points surpassed the corresponding thresholds was considered a significant cluster. The minimum duration (T) of a significant cluster was tested from 20 ms to 100 ms with a step size of 10 ms to observe the impact of T on the detected onset times and the relative time lag among different features in each task. Generally, when T was set to 20, 30, and 40 ms, the onset times of all features were within 100 ms, which was consistent with previous research (Teichmann et al., 2019, 2020, 2021; Cichy et al., 2014; Cichy & Pantazis, 2017; Ritchie et al., 2015). When T was adjusted to more than 40 ms, the onset times of features exceeded 100 ms. As T increased, the onset times were delayed, and the patterns of the time lags among features, although obscured, remained (examples in Figure S5). Previous empirical research found early onset latencies of decoding accuracy for various categorization levels to be between 48-70 ms (Cichy et al., 2014; Ritchie et al., 2015b). Among the various values of T we tested, our observed onset times were closest to those reported in previous studies when T was set to 20 ms. Therefore, we determined that a time window of at least 20 ms, where all time points exceeded the corresponding threshold, was considered a significant cluster containing feature information. In each trial, the first time point of the first significant cluster was considered the earliest time point at which the feature was represented. The time point with the highest d-value across all significant clusters in each trial was considered the peak time point. It should be noted that the peak time within the reaction time of each trial was considered effective and used for further analysis. Ultimately, for each participant, ROI, and task type, we obtained onset and peak times for color, animacy, and size features for each trial.

### GLMM analyses

Following Linde-Domingo et al., 2019, we employed generalized linear mixed models (GLMMs) to examine our hypotheses regarding the relative onset or peak time lags of d-values among different features within the same categorization task during the perception and retrieval stages. We chose GLMMs over the more commonly used GLM-based models (i.e., ANOVAs or t-tests) because they make fewer assumptions about the data distribution and are better suited for modeling d-value onsets. The task type (color, animacy and size tasks), feature type (color, animacy and size features), and their interactions were modeled as fixed effects in the GLMM. By examining the interaction between task type and feature type, we assessed whether selective attention modulated the onset or peak times of different features. Participant ID (including the intercept) was modeled as a random factor. All models for analyzing d-value onset or peak times used a gamma probability distribution and an identity link function, as employed by Linde-Domingo et al., 2019.

### Phase coupling estimation

If selective attention can modulate the times of feature representations, how is this process achieved? We hypothesized that during the fixation period (i.e., the task preparation stage) when stimuli (or spatial cues) were not presented, the brain networks were configured to be prepared for the appropriate execution of the instructed task. The stronger the connectivity between brain regions, the more prepared the configured brain networks are, and the faster the target features might be processed. We used the phase slope index (PSI) to characterize the connectivity patterns between the occipital and the other three higher-order regions (the temporal, parietal, and frontal lobes) and between the temporal and frontal lobes (a control analysis to contrast with the retrieval task) during the preparation stage in the feature perception task. For the preparation stage in the feature retrieval task, we calculated the PSI between the temporal and frontal lobes. The PSI is a method to estimate the phase coupling patterns between brain regions and to detect the direction of frequency-specific neural interactions from time series, determining which of the two phase-coupled brain areas is the ‘leader’ (Nolte et al., 2008). Compared to Granger causality, the phase slope index does not yield significant false detections for mixtures of independent sources (Nolte et al., 2008). The PSI between two time series for channels *i* and *j* is defined as

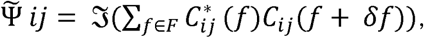

where 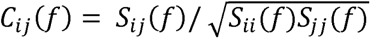 is the complex coherency, *S* is the cross-spectral matrix, δf is the frequency resolution, and 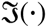 denotes taking the imaginary part. *F* is the set of frequencies over which the slope is summed. We used the continuous Morlet wavelet transform to estimate the cross-spectral matrix.

Before calculating PSI, we employed principal component analysis (PCA), also known as spatial filters, to transform the channel data to new sources or virtual channels for each participant and ROI. This step aimed to reduce computation time and increase the signal-to-noise ratio (SNR). The PCA analysis was conducted separately for the preparation stage in the feature perception and retrieval tasks. The representative time courses in each ROI were defined as the smallest set of signals that explained at least 90% of the raw amplitude of the pre-processed MEG time courses.

PSI values were calculated for all pairs of virtual channels of each pair of ROIs with a frequency resolution of 1 Hz for each frequency band (theta: 4-8 Hz; alpha: 9-13 Hz; beta: 14-30 Hz; gamma: 31-100 Hz). For each pair of ROIs, the higher-order ROIs were set as seed ROIs, and the occipital lobe was set as the target ROI. For example, participant 1 had n virtual channels in the temporal lobe (the seed ROI) and m virtual channels in the occipital lobe (the target ROI), the PSI values for the temporal-occipital pair would form an n×m matrix for each time point. The sign of PSI values indicates the direction of information transfer. For instance, when the temporal and occipital cortex were set as the seed and target ROI, respectively, a positive PSI value means the information transfer direction is from the temporal to the occipital lobe, while a negative PSI value indicates the opposite direction. We employed cluster-based permutation analyses to investigate in which time ranges the PSI values of each task, frequency band, and pair of ROIs differed significantly from zero. This analysis indicated the presence of directional phase coupling between ROIs, helping us understand the neural connectivity patterns that underpin selective attention modulation during the task preparation stages.

After identifying the significant clusters of PSI at the group level for the preparation stage in the feature perception task, we extracted the PSI values from the corresponding time range of significant clusters for each participant and averaged them across the time points in each cluster. When the PSI values of a significant cluster were negative, they were converted to positive values, with larger values indicating stronger phase coupling. If there were more than one significant cluster in a condition, the PSI values were averaged across clusters. We then performed a Pearson’s correlation between the averaged PSI values and the onset times of features in the target ROI to investigate whether stronger phase coupling between brain regions during the task preparation stage correlated with earlier onset times of target features instead of non-target features during the perception stage. For each participant and ROI, the onset time of target features was calculated as the average of the onset times of the color feature in the color task, the animacy feature in the animacy task, and the size feature in the size task. The onset time of non-target features was calculated as the average of the onset times of the color feature in the animacy and size tasks, the animacy feature in the color and size tasks, and the size feature in the color and animacy tasks.

## Acknowledgements

This work was supported by the National Natural Science Foundation of China (32330039) and the Fundamental Research Funds for the Central Universities (2243300006)

## Supplementary materials

**Figure S1.**
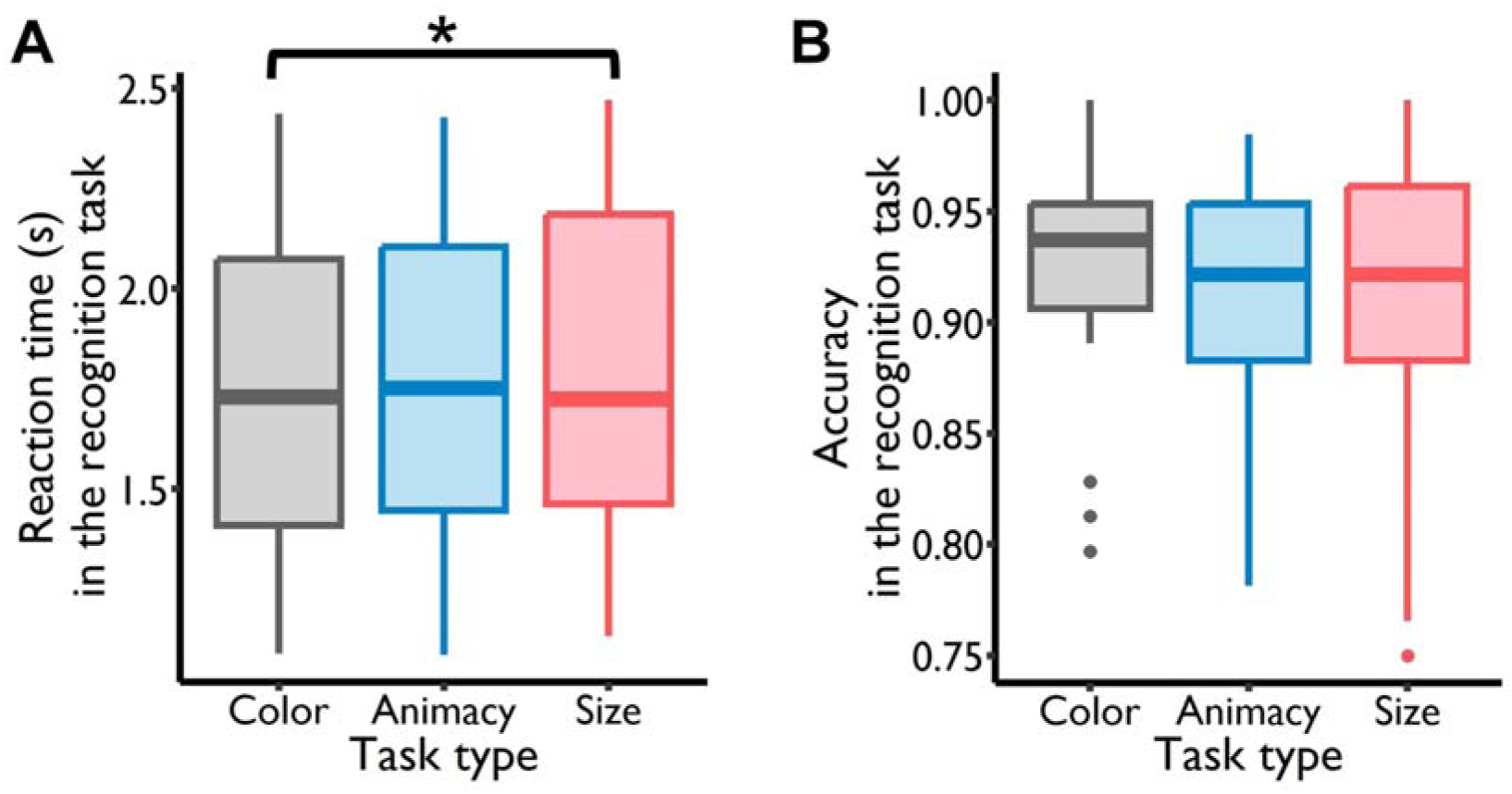
The behavioral results in the recognition task. **A. Reaction time.** The mean reaction times of the object recognition under the color, animacy and size task conditions were 1.732 s (*SD* = 0.403 s), 1.766 s (*SD* = 0.402 s), and 1.822 s (*SD* = 0.412 s). The task type predicted the reaction times (*F_2, 52_* = 4.255, *p* = 0.019). Post-hoc testing showed that the reaction time of the object recognition under the color task condition was significantly faster than that under the size task condition (*t_52_* = -2.889, *p_corrected_* = 0.015), while the difference between the animacy task condition and the other two task conditions was not significant (animacy condition vs. color condition: *t_52_* = 1.097, *p_corrected_*= 0.520; animacy condition vs. size condition: *t_52_* = -1.793, *p_corrected_* = 0.182). **B. Accuracy.** The mean accuracy of the object recognition under the color, animacy and size task conditions was 92.59% (*SD* = 5.44%), 91.38% (*SD* = 5.12%) and 91.55% (*SD* = 6.21%). The task type didn’t predict the accuracy of object recognition under three task conditions (*F_2, 52_*= 1.181, *p* = 0.315).

**Figure S2.**
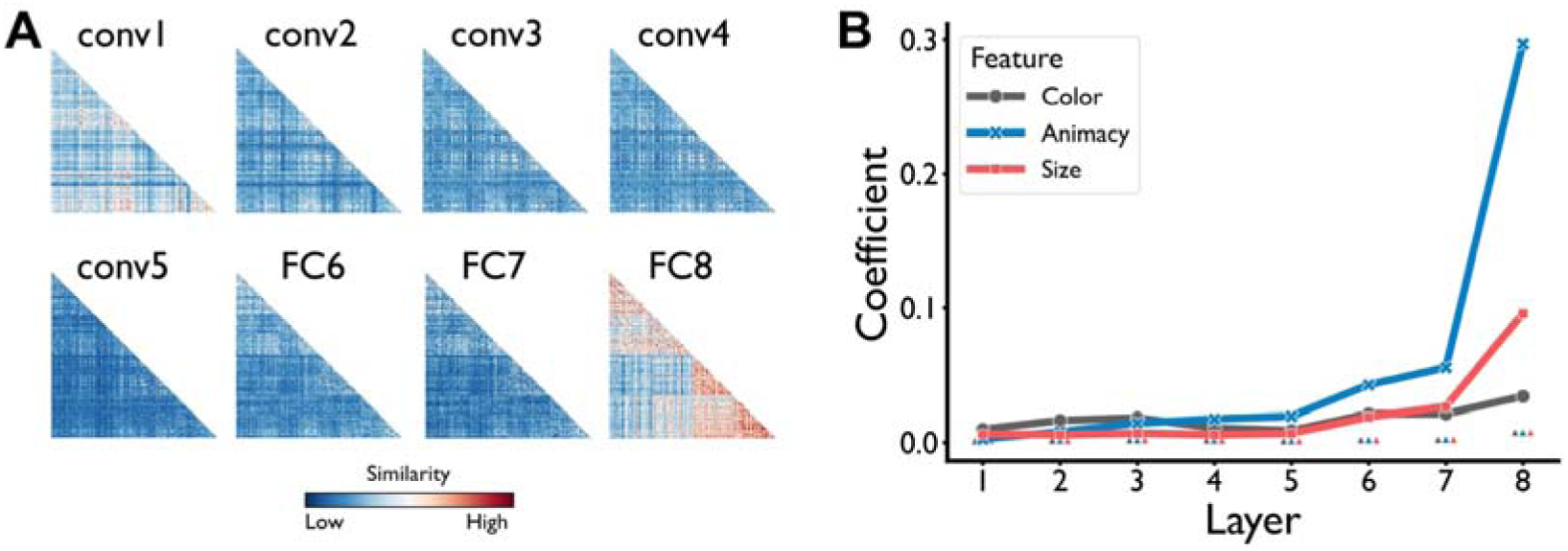
Representations of three features (color, animacy and size) in the deep neural network AlexNet (stimulus set 2). **A. Representational similarity matrix.** The matrix was generated by correlating the activations of the artificial neurons in each DNN layer. **B. Sensitivity of feature information.** The threshold of feature-specific representations in each layer was indicated by the little triangles which were not connected by lines. This figure was generated by the data of the stimulus set 2. conv: convolutional layer; FC: fully connected layer.

**Figure S3.**
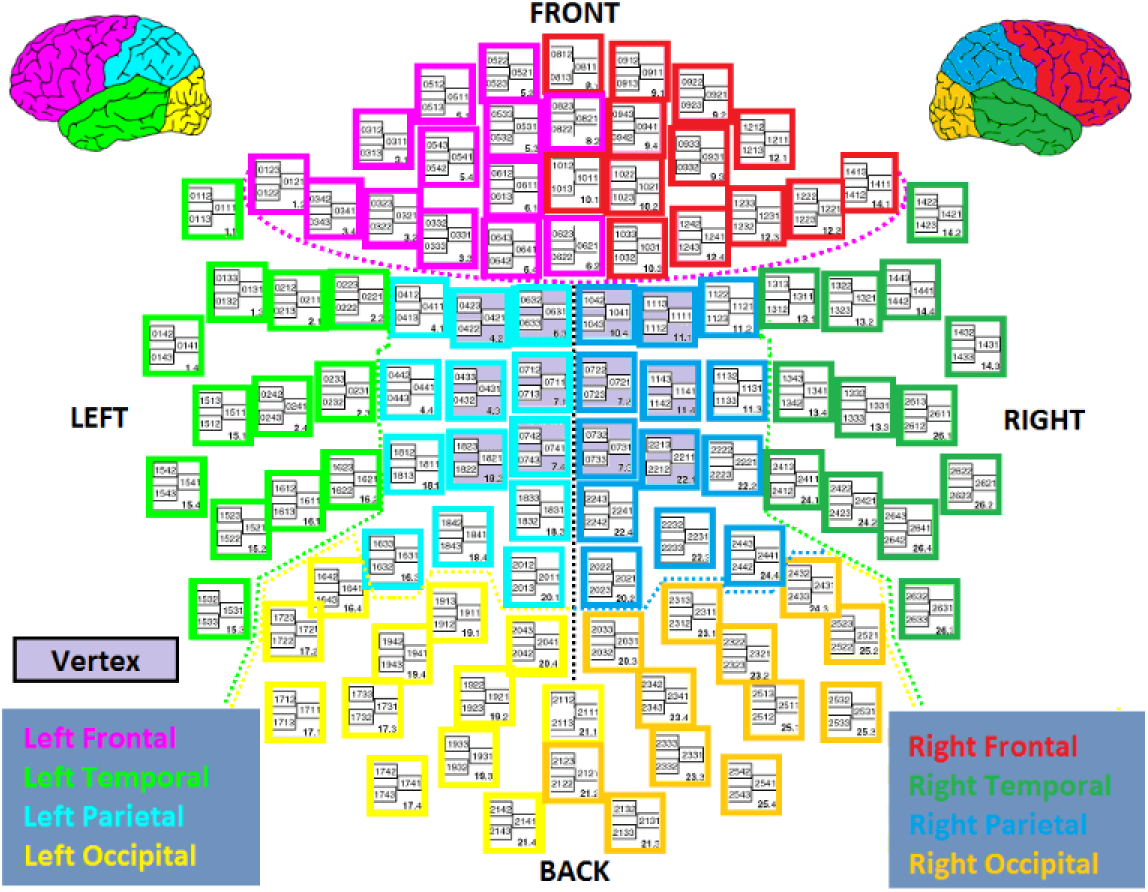
Elekta Neuromag MEG channel positions. Channels corresponding to different lobes are color-coded (figure adapted from www.megwiki.org)

**Figure S4.**
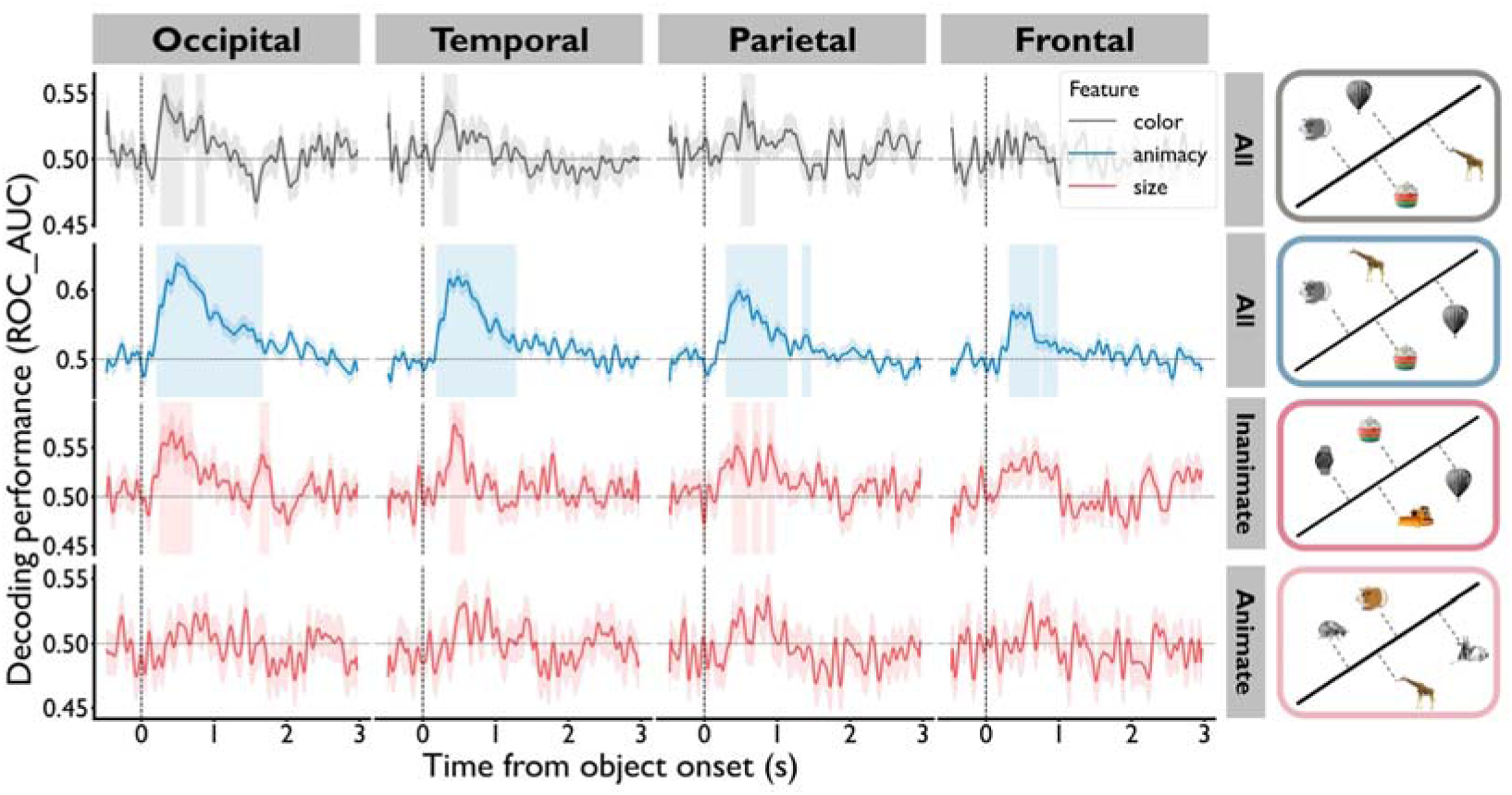
The decoding performance of three features during the learning stage. Significant clusters of color, animacy and size features in the occipital, temporal and parietal cortex were found (*P_corrected_* < 0.05). In the frontal cortex, only significant clusters of the animacy feature were found (*P_corrected_* < 0.05). There were no significant clusters of size feature for the animate objects. The shaded areas surrounding the classification performance time courses indicated standard error across participants. The vertical shaded areas marked the significant clusters carrying the feature information.

**Figure S5.**
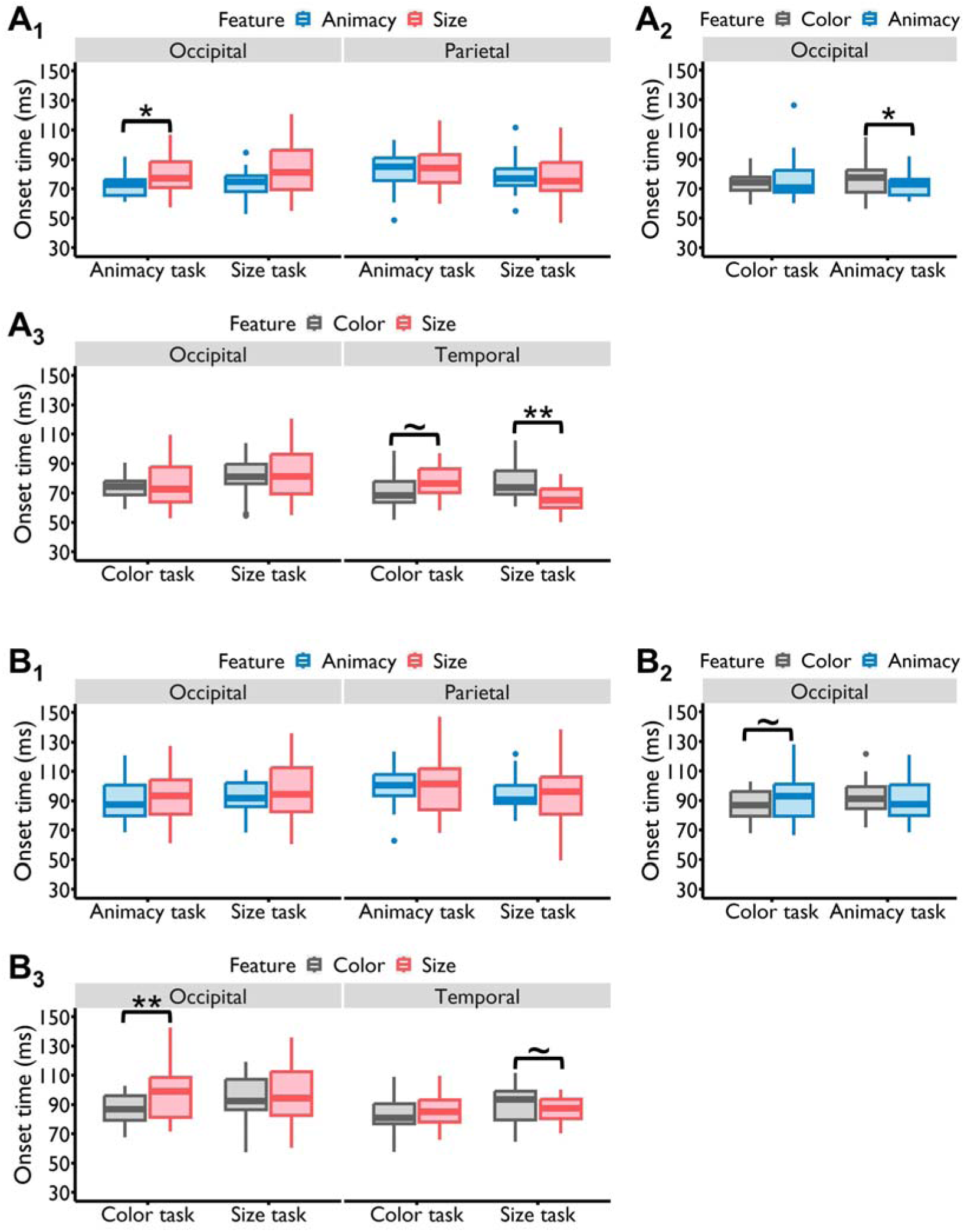
The modulation effects of selective attention when setting T=30 ms. **(A) or 40 ms (B)**, i.e., the time window of at least 30 ms or 40 ms in which the d-values of all time points exceeding the corresponding threshold was considered as a significant cluster containing feature information. As shown in the figures, when the minimum duration (T) of a significant cluster increased, the overall onset times of features were delayed accordingly, and the patterns of time lags between different features remained discernible, although some of them became obscured.

**Figure S6.**
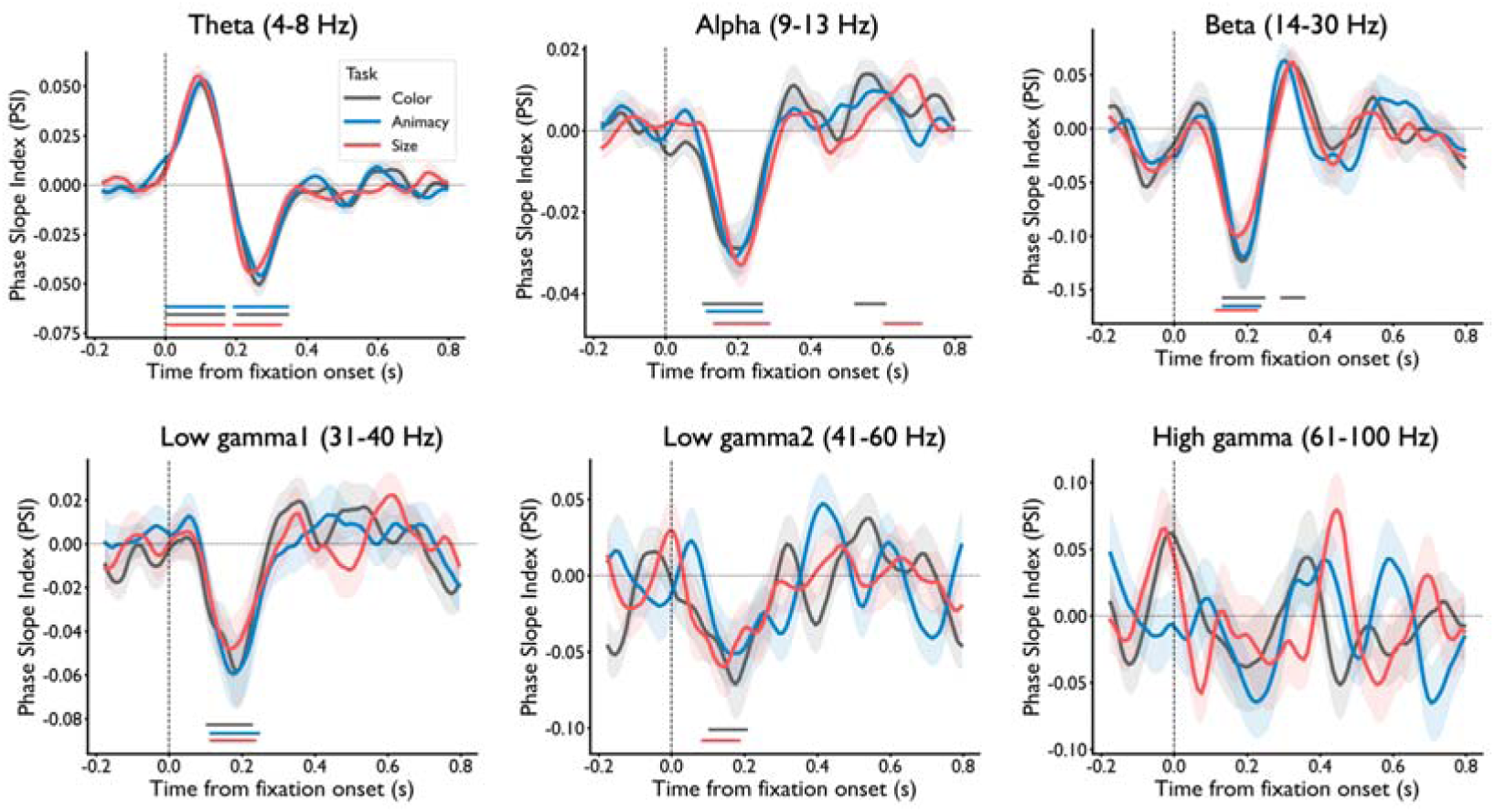
Phase coupling patterns between the temporal and occipital lobes across different frequency bands. Only phase coupling in the theta band demonstrated a pattern of initially top-down followed by bottom-up interactions. In contrast, the phase couplings in the alpha, beta, and low gamma bands were predominantly bottom-up, indicating an information flow from the occipital to temporal lobes. No significant phase coupling was observed in the high gamma band.

**Table S1.**
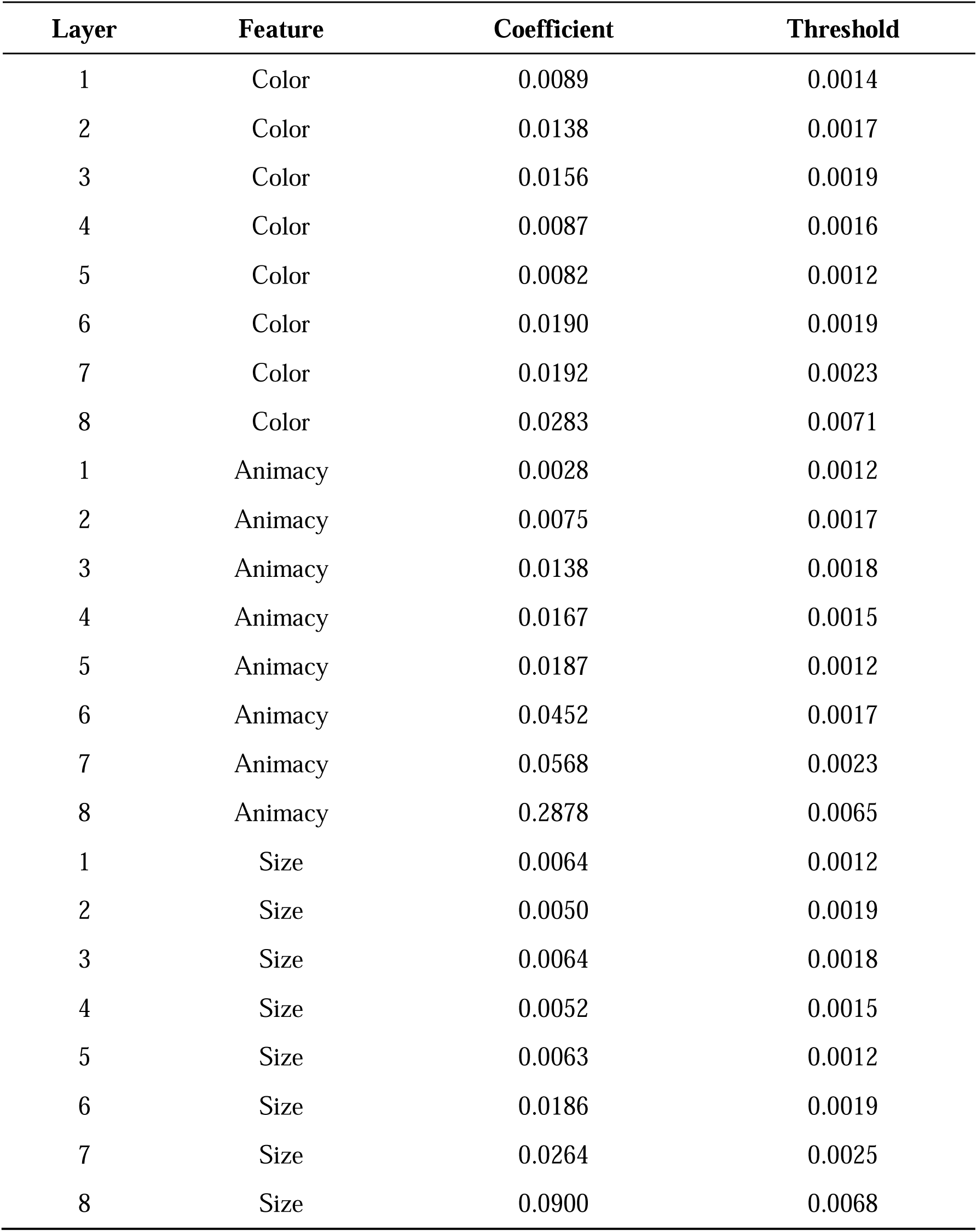
The coefficients of features for stimulus set 1.

**Table S2.**
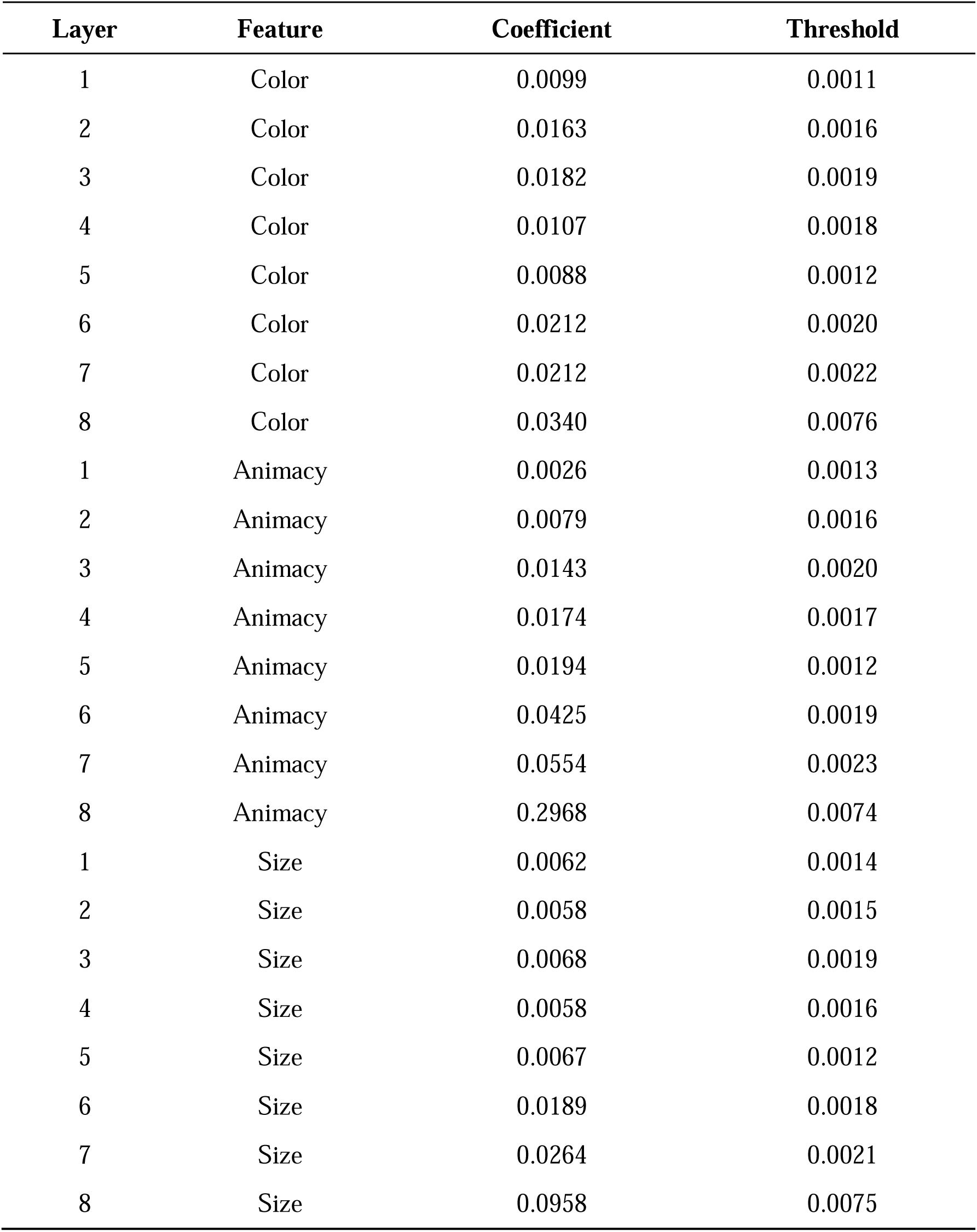
The coefficients of features for stimulus set 2.

## References

1. Baldauf, D., & Desimone, R. (2014). Neural Mechanisms of Object-Based Attention. Science, 344(6182), 424–427. 10.1126/science.1247003

2. Bone, M. B., Ahmad, F., & Buchsbaum, B. R. (2020). Feature-specific neural reactivation during episodic memory. Nature Communications, 11(1), 1945. 10.1038/s41467-020-15763-2

3. Bouchacourt, F., & Buschman, T. J. (2019). A Flexible Model of Working Memory. Neuron, 103(1), 147–160.e8. 10.1016/j.neuron.2019.04.020

4. Brincat, S. L., & Connor, C. E. (2004). Underlying principles of visual shape selectivity in posterior inferotemporal cortex. Nature Neuroscience, 7(8), 880–886. 10.1038/nn1278

5. Bürkner, P.-C. (2017). brms: An R Package for Bayesian Multilevel Models Using Stan. Journal of Statistical Software, 80(1), 1–28. 10.18637/jss.v080.i01

6. Buschman, T. J., & Kastner, S. (2015). From Behavior to Neural Dynamics: An Integrated Theory of Attention. Neuron, 88(1), 127–144. 10.1016/j.neuron.2015.09.017

7. Carlson, T., Tovar, D. A., Alink, A., & Kriegeskorte, N. (2013). Representational dynamics of object vision: The first 1000 ms. Journal of Vision, 13(10), 1–1. 10.1167/13.10.1

8. Chatham, C. H., Frank, M. J., & Badre, D. (2014). Corticostriatal Output Gating during Selection from Working Memory. Neuron, 81(4), 930–942. 10.1016/j.neuron.2014.01.002

9. Cichy, R. M., & Pantazis, D. (2017). Multivariate pattern analysis of MEG and EEG: A comparison of representational structure in time and space. NeuroImage, 158(July), 441–454. 10.1016/j.neuroimage.2017.07.023

10. Cichy, R. M., Pantazis, D., & Oliva, A. (2014). Resolving human object recognition in space and time. Nature Neuroscience, 17(3), 455–462. 10.1038/nn.3635

11. Clarke, A., Taylor, K. I., Devereux, B., Randall, B., & Tyler, L. K. (2013). From Perception to Conception: How Meaningful Objects Are Processed over Time. Cerebral Cortex, 23(1), 187–197. 10.1093/cercor/bhs002

12. Córdova, N. I., Tompary, A., & Turk-Browne, N. B. (2016). Attentional modulation of background connectivity between ventral visual cortex and the medial temporal lobe. Neurobiology of Learning and Memory, 134, 115–122. 10.1016/j.nlm.2016.06.011

13. Deng, J., Dong, W., Socher, R., Li, L.-J., Kai Li, & Li Fei-Fei. (2009). ImageNet: A large-scale hierarchical image database. 2009 IEEE Conference on Computer Vision and Pattern Recognition, 248–255. 10.1109/CVPR.2009.5206848

14. Desimone, R., & Duncan, J. (1995). NEURAL MECHANISMS OF SELECTIVE VISUAL ATTENTION. Annu. Rev. Neurosci, 18, 193–222.

15. Egner, T., & Hirsch, J. (2005). Cognitive control mechanisms resolve conflict through cortical amplification of task-relevant information. Nature Neuroscience, 8(12), 1784–1790. 10.1038/nn1594

16. Engel, A. K., Fries, P., & Singer, W. (2001). Dynamic predictions: Oscillations and synchrony in top–down processing. Nature Reviews Neuroscience, 2(10), 704–716. 10.1038/35094565

17. Ester, E. F., Sutterer, D. W., Serences, J. T., & Awh, E. (2016). Feature-Selective Attentional Modulations in Human Frontoparietal Cortex. Journal of Neuroscience, 36(31), 8188–8199. 10.1523/JNEUROSCI.3935-15.2016

18. Firestone, C., & Scholl, B. J. (2016). Cognition does not affect perception: Evaluating the evidence for “top-down” effects. Behavioral and Brain Sciences, 39, e229. 10.1017/S0140525X15000965

19. Fuster, J. M., Bauer, R. H., & Jervey, J. P. (1985). Functional Interactions Between Inferotemporal and Prefrontal Cortex in a Cognitive Task. Brain Research, 330(1985), 299–307.

20. Gazzaley, A., & Nobre, A. C. (2012). Top-down modulation: Bridging selective attention and working memory. Trends in Cognitive Sciences, 16(2), 129–135. 10.1016/j.tics.2011.11.014

21. Gerbella, M., Belmalih, A., Borra, E., Rozzi, S., & Luppino, G. (2010). Cortical Connections of the Macaque Caudal Ventrolateral Prefrontal Areas 45A and 45B. Cerebral Cortex, 20(1), 141–168. 10.1093/cercor/bhp087

22. Gibson, J. J. (1950). The Perception Of The Visual World (Vol. 60, Issue 4, pp. 594–595). Houghton Mifflin.

23. Gramfort, A., Luessi, M., Larson, E., Engemann, D. A., Strohmeier, D., Brodbeck, C., Goj, R., Jas, M., Brooks, T., Parkkonen, L., & Hämäläinen, M. (2013). MEG and EEG data analysis with MNE-Python. Frontiers in Neuroscience, *7*(7 DEC), 1–13. 10.3389/fnins.2013.00267

24. Grootswagers, T., Robinson, A. K., Shatek, S. M., & Carlson, T. A. (2021). The neural dynamics underlying prioritisation of task-relevant information. *Neurons, Behavior*, Data Analysis, and Theory, 5(1). 10.51628/001c.21174

25. Hebart, M. N., & Baker, C. I. (2018). Deconstructing multivariate decoding for the study of brain function. NeuroImage, 180, 4–18. 10.1016/j.neuroimage.2017.08.005

26. Jas, M., Engemann, D. A., Bekhti, Y., Raimondo, F., & Gramfort, A. (2017). Autoreject: Automated artifact rejection for MEG and EEG data. NeuroImage, 159(December 2016), 417–429. 10.1016/j.neuroimage.2017.06.030

27. Jehee, J. F. M., Brady, D. K., & Tong, F. (2011). Attention Improves Encoding of Task-Relevant Features in the Human Visual Cortex. The Journal of Neuroscience, 31(22), 8210–8219. 10.1523/JNEUROSCI.6153-09.2011

28. Khaligh-Razavi, S.-M., Cichy, R. M., Pantazis, D., & Oliva, A. (2018). Tracking the Spatiotemporal Neural Dynamics of Real-world Object Size and Animacy in the Human Brain. Journal of Cognitive Neuroscience, 30(11), 1559–1576. 10.1162/jocn_a_01290

29. King, J. R., Pescetelli, N., & Dehaene, S. (2016). Brain Mechanisms Underlying the Brief Maintenance of Seen and Unseen Sensory Information. Neuron, 92(5), 1122–1134. 10.1016/j.neuron.2016.10.051

30. Konkle, T., & Caramazza, A. (2013). Tripartite organization of the ventral stream by animacy and object size. Journal of Neuroscience, 33(25), 10235–10242. 10.1523/JNEUROSCI.0983-13.2013

31. Kravitz, D. J., Saleem, K. S., Baker, C. I., Ungerleider, L. G., & Mishkin, M. (2013). The ventral visual pathway: An expanded neural framework for the processing of object quality. Trends in Cognitive Sciences, 17(1), 26–49. 10.1016/j.tics.2012.10.011

32. Kriegeskorte, N., Goebel, R., & Bandettini, P. (2006). Information-based functional brain mapping. Proceedings of the National Academy of Sciences, 103(10), 3863–3868. 10.1073/pnas.0600244103

33. Krizhevsky, A., Sutskever, I., & Hinton, G. E. (2017). ImageNet classification with deep convolutional neural networks. Communications of the ACM, 60(6), 84–90. 10.1145/3065386

34. Linde-Domingo, J., Treder, M. S., Kerrén, C., & Wimber, M. (2019). Evidence that neural information flow is reversed between object perception and object reconstruction from memory. Nature Communications, 10(1), 179. 10.1038/s41467-018-08080-2

35. Liu, Y., Dolan, R. J., Kurth-Nelson, Z., & Behrens, T. E. J. (2019). Human Replay Spontaneously Reorganizes Experience. Cell, 178(3), 640–652.e14. 10.1016/j.cell.2019.06.012

36. Long, N. M., & Kuhl, B. A. (2018). Bottom-Up and Top-Down Factors Differentially Influence Stimulus Representations Across Large-Scale Attentional Networks. The Journal of Neuroscience, 38(10), 2495–2504. 10.1523/JNEUROSCI.2724-17.2018

37. Lu, Y., Wang, C., Chen, C., & Xue, G. (2015). Spatiotemporal Neural Pattern Similarity Supports Episodic Memory. Current Biology, 25(6), 780–785. 10.1016/j.cub.2015.01.055

38. Mack, M. L., & Palmeri, T. J. (2011). The Timing of Visual Object Categorization. Frontiers in Psychology, 2. 10.3389/fpsyg.2011.00165

39. Mante, V., Sussillo, D., Shenoy, K. V., & Newsome, W. T. (2013). Context-dependent computation by recurrent dynamics in prefrontal cortex. Nature, 503(7474), 78–84. 10.1038/nature12742

40. Maris, E., & Oostenveld, R. (2007). Nonparametric statistical testing of EEG- and MEG-data. Journal of Neuroscience Methods, 164(1), 177–190. 10.1016/j.jneumeth.2007.03.024

41. Maunsell, J. H. R., & Treue, S. (2006). Feature-based attention in visual cortex. Trends in Neurosciences, 29(6), 317–322. 10.1016/j.tins.2006.04.001

42. Mirjalili, S., Powell, P., Strunk, J., James, T., & Duarte, A. (2021). Context Memory Encoding and Retrieval Temporal Dynamics are Modulated by Attention across the Adult Lifespan. Eneuro, 8(1), ENEURO.0387-20.2020. 10.1523/ENEURO.0387-20.2020

43. Moore, T., & Zirnsak, M. (2017). Neural Mechanisms of Selective Visual Attention. Annual Review of Psychology, 68(1), 47–72. 10.1146/annurev-psych-122414-033400

44. 44. Nastase, S. A., Connolly, A. C., Oosterhof, N. N., Halchenko, Y. O., Guntupalli, J. S., Visconti di Oleggio Castello, M., Gors, J., Gobbini, M. I., & Haxby, J. V. (2017). Attention Selectively Reshapes the Geometry of Distributed Semantic Representation. Cerebral Cortex, 27(8), 4277–4291. 10.1093/cercor/bhx138

45. Nee, D. E., & Jonides, J. (2008). Neural correlates of access to short-term memory. Proceedings of the National Academy of Sciences, 105(37), 14228–14233. 10.1073/pnas.0802081105

46. Nobre, A. C., Coull, J. T., Maquet, P., Frith, C. D., Vandenberghe, R., & Mesulam, M. M. (2004). Orienting Attention to Locations in Perceptual Versus Mental Representations. Journal of Cognitive Neuroscience, 16(3), 363–373. 10.1162/089892904322926700

47. 47. Nobre, A. C. (Kia), & Kastner, S. (Eds.). (2014). The Oxford Handbook of Attention. Oxford University Press. 10.1093/oxfordhb/9780199675111.001.0001

48. Nolte, G., Ziehe, A., Nikulin, V. V., Schlögl, A., Krämer, N., Brismar, T., & Müller, K.-R. (2008). Robustly Estimating the Flow Direction of Information in Complex Physical Systems. Physical Review Letters, 100(23), 234101. 10.1103/PhysRevLett.100.234101

49. O’Regan, J. K., & Noë, A. (2001). A sensorimotor account of vision and visual consciousness. Behavioral and Brain Sciences, 24(5), 939–973. 10.1017/S0140525X01000115

50. Peirce, J., Gray, J. R., Simpson, S., MacAskill, M., Höchenberger, R., Sogo, H., Kastman, E., & Lindeløv, J. K. (2019). PsychoPy2: Experiments in behavior made easy. Behavior Research Methods, 51(1), 195–203. 10.3758/s13428-018-01193-y

51. Ramezanpour, H., & Fallah, M. (2022). The role of temporal cortex in the control of attention. Current Research in Neurobiology, 3, 100038. 10.1016/j.crneur.2022.100038

52. Ratcliff, R. (1978). A theory of memory retrieval. Psychological Review, 85(2), 59–108. 10.1037/0033-295X.85.2.59

53. Ratcliff, R., & McKoon, G. (2008a). The Diffusion Decision Model: Theory and Data for Two-Choice Decision Tasks. Neural Computation, 20(4), 873–922. 10.1162/neco.2008.12-06-420

54. Ratcliff, R., & McKoon, G. (2008b). The Diffusion Decision Model: Theory and Data for Two-Choice Decision Tasks. Neural Computation, 20(4), 873–922. 10.1162/neco.2008.12-06-420

55. Ritchie, J. B., Tovar, D. A., & Carlson, T. A. (2015a). Emerging Object Representations in the Visual System Predict Reaction Times for Categorization. PLOS Computational Biology, 11(6), e1004316. 10.1371/journal.pcbi.1004316

56. Ritchie, J. B., Tovar, D. A., & Carlson, T. A. (2015b). Emerging Object Representations in the Visual System Predict Reaction Times for Categorization. PLoS Computational Biology, 11(6), 1–19. 10.1371/journal.pcbi.1004316

57. Sakai, K. (2008). Task Set and Prefrontal Cortex. Annual Review of Neuroscience, 31(1), 219–245. 10.1146/annurev.neuro.31.060407.125642

58. Saleem, K. S., Miller, B., & Price, J. L. (2014). Subdivisions and connectional networks of the lateral prefrontal cortex in the macaque monkey. Journal of Comparative Neurology, 522(7), 1641–1690. 10.1002/cne.23498

59. Serences, J. T., & Yantis, S. (2006). Selective visual attention and perceptual coherence. Trends in Cognitive Sciences, 10(1), 38–45. 10.1016/j.tics.2005.11.008

60. Serre, T., Oliva, A., & Poggio, T. (2007). A feedforward architecture accounts for rapid categorization. Proceedings of the National Academy of Sciences, 104(15), 6424–6429. 10.1073/pnas.0700622104

61. Sheng, J., Zhang, L., Liu, C., Liu, J., Feng, J., Zhou, Y., Hu, H., & Xue, G. (2022). Higher-dimensional neural representations predict better episodic memory. Science Advances, 8(16), eabm3829. 10.1126/sciadv.abm3829

62. Siegel, M., Donner, T. H., Oostenveld, R., Fries, P., & Engel, A. K. (2008). Neuronal Synchronization along the Dorsal Visual Pathway Reflects the Focus of Spatial Attention. Neuron, 60(4), 709–719. 10.1016/j.neuron.2008.09.010

63. Sprague, T. C., & Serences, J. T. (2013). Attention modulates spatial priority maps in the human occipital, parietal and frontal cortices. Nature Neuroscience, 16(12), 1879–1887. 10.1038/nn.3574

64. Stone, M. (1960). Models for choice-reaction time. Psychometrika, 25(3), 251–260. 10.1007/BF02289729

65. 65. Summerfield, C., & De Lange, F. P. (2014). Expectation in perceptual decision making: Neural and computational mechanisms. Nature Reviews Neuroscience, 15(11), 745–756. 10.1038/nrn3838

66. Taulu, S., & Simola, J. (2006). Spatiotemporal signal space separation method for rejecting nearby interference in MEG measurements. Physics in Medicine and Biology, 51(7), 1759–1768. 10.1088/0031-9155/51/7/008

67. Teichmann, L., Grootswagers, T., Carlson, T. A., & Rich, A. N. (2019). Seeing versus knowing: The temporal dynamics of real and implied colour processing in the human brain. NeuroImage, 200(October 2018), 373–381. 10.1016/j.neuroimage.2019.06.062

68. Teichmann, L., Grootswagers, T., Moerel, D., Carlson, T. A., & Rich, A. N. (2021). Temporal dissociation of neural activity underlying synesthetic and perceptual colors. Proceedings of the National Academy of Sciences of the United States of America, 118(6), 2–4. 10.1073/pnas.2020434118

69. Teichmann, L., Quek, G. L., Robinson, A. K., Grootswagers, T., Carlson, T. A., & Rich, A. N. (2020). Yellow strawberries and red bananas: The influence of object-colour knowledge on emerging object representations in the brain. The Journal of Neuroscience, 40(35), 6779–6789. 10.1101/533513

70. Tijl Grootswagers, Susan G. Wardle, and T. A. C. (2017). Decoding Dynamic Brain Patterns from Evoked Responses: A Tutorial on Multivariate Pattern Analysis Applied to Time Series Neuroimaging Data. Journal of Cognitive Neuroscience, 1–10. 10.1162/jocn

71. Tomita, H., Ohbayashi, M., Nakahara, K., Hasegawa, I., & Miyashita, Y. (1999). Top-down signal from prefrontal cortex in executive control of memory retrieval. Nature, 401(6754), 699–703. 10.1038/44372

72. Treisman, A. M. (1980). A Feature-Integration Theory of Attention. Cognitive Psychology, 12(1), 97–136.

73. Wang, R., Janini, D., & Konkle, T. (2022a). Mid-level Feature Differences Support Early Animacy and Object Size Distinctions: Evidence from Electroencephalography Decoding. Journal of Cognitive Neuroscience, 34(9), 1670–1680. 10.1162/jocn_a_01883

74. Wang, R., Janini, D., & Konkle, T. (2022b). Mid-level feature differences underlie early animacy and object size distinctions: Evidence from EEG decoding. bioRxiv, 2022.01.12.475180. 10.1101/2022.01.12.475180

75. Wimmer, G. E., Liu, Y., Vehar, N., Behrens, T. E. J., & Dolan, R. J. (2020). Episodic memory retrieval success is associated with rapid replay of episode content. Nature Neuroscience, 23(8), 1025–1033. 10.1038/s41593-020-0649-z

76. Xue, G., Dong, Q., Chen, C., Lu, Z.-L., Mumford, J. A., & Poldrack, R. A. (2013). Complementary Role of Frontoparietal Activity and Cortical Pattern Similarity in Successful Episodic Memory Encoding. Cerebral Cortex, 23(7), 1562–1571. 10.1093/cercor/bhs143

77. Yau, J. M., Pasupathy, A., Brincat, S. L., & Connor, C. E. (2013). Curvature Processing Dynamics in Macaque Area V4. Cerebral Cortex, 23(1), 198–209. 10.1093/cercor/bhs004

78. Yeo, B. T. T., Krienen, F. M., Sepulcre, J., Sabuncu, M. R., Lashkari, D., Hollinshead, M., Roffman, J. L., Smoller, J. W., Zöllei, L., Polimeni, J. R., Fischl, B., Liu, H., & Buckner, R. L. (2011). The organization of the human cerebral cortex estimated by intrinsic functional connectivity. J Neurophysiol, 106.

79. Zheng, L., Gao, Z., Xiao, X., Ye, Z., Chen, C., & Xue, G. (2018). Reduced Fidelity of Neural Representation Underlies Episodic Memory Decline in Normal Aging. Cerebral Cortex, 28(7), 2283–2296. 10.1093/cercor/bhx130

80. Zhou, T., Kawasaki, K., Suzuki, T., Hasegawa, I., Roe, A. W., & Tanigawa, H. (2023). Mapping information flow between the inferotemporal and prefrontal cortices via neural oscillations in memory retrieval and maintenance. Cell Reports, 42(10), 113169. 10.1016/j.celrep.2023.113169

