## Supplemental Figures and Tables for "Dynamic Modulation of Representational Trajectories Through Selective Attention"

### Supplementary materials

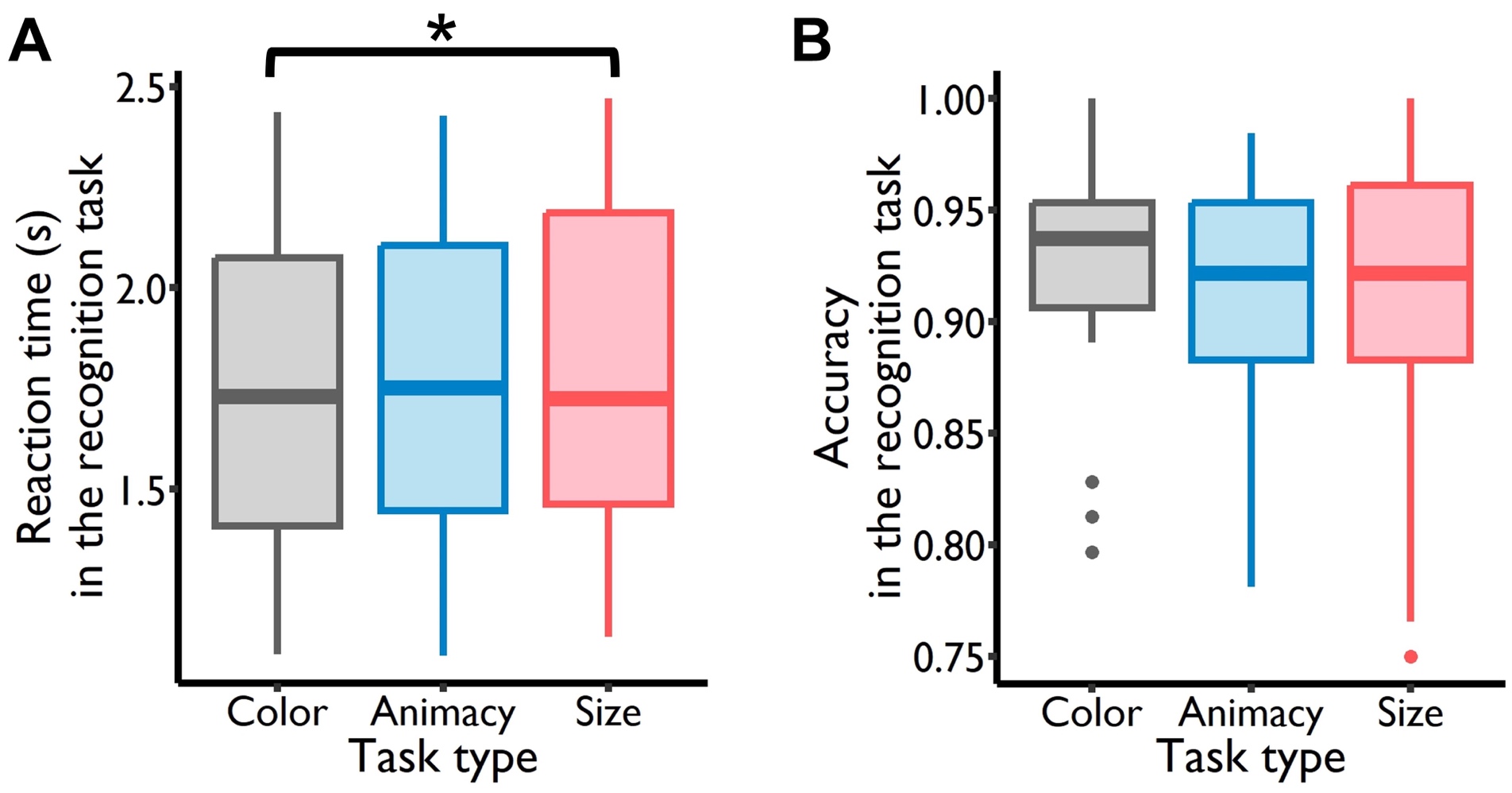

**Figure S1. The behavioral results in the recognition task. A. Reaction time.** The mean reaction times of the object recognition under the color, animacy and size task conditions were 1.732 s (*SD* = 0.403 s), 1.766 s (*SD* = 0.402 s), and 1.822 s (*SD* = 0.412 s). The task type predicted the reaction times (*F_2, 52_* = 4.255, *p* = 0.019). Post-hoc testing showed that the reaction time of the object recognition under the color task condition was significantly faster than that under the size task condition (*t_52_* = -2.889, *p_corrected_* = 0.015), while the difference between the animacy task condition and the other two task conditions was not significant (animacy condition vs. color condition: *t_52_* = 1.097, *p_corrected_* = 0.520; animacy condition vs. size condition: *t_52_* = -1.793, *p_corrected_* = 0.182). **B. Accuracy.** The mean accuracy of the object recognition under the color, animacy and size task conditions was 92.59% (*SD* = 5.44%), 91.38% (*SD* = 5.12%) and 91.55% (*SD* = 6.21%). The task type didn’t predict the accuracy of object recognition under three task conditions (*F_2, 52_* = 1.181, *p* = 0.315).

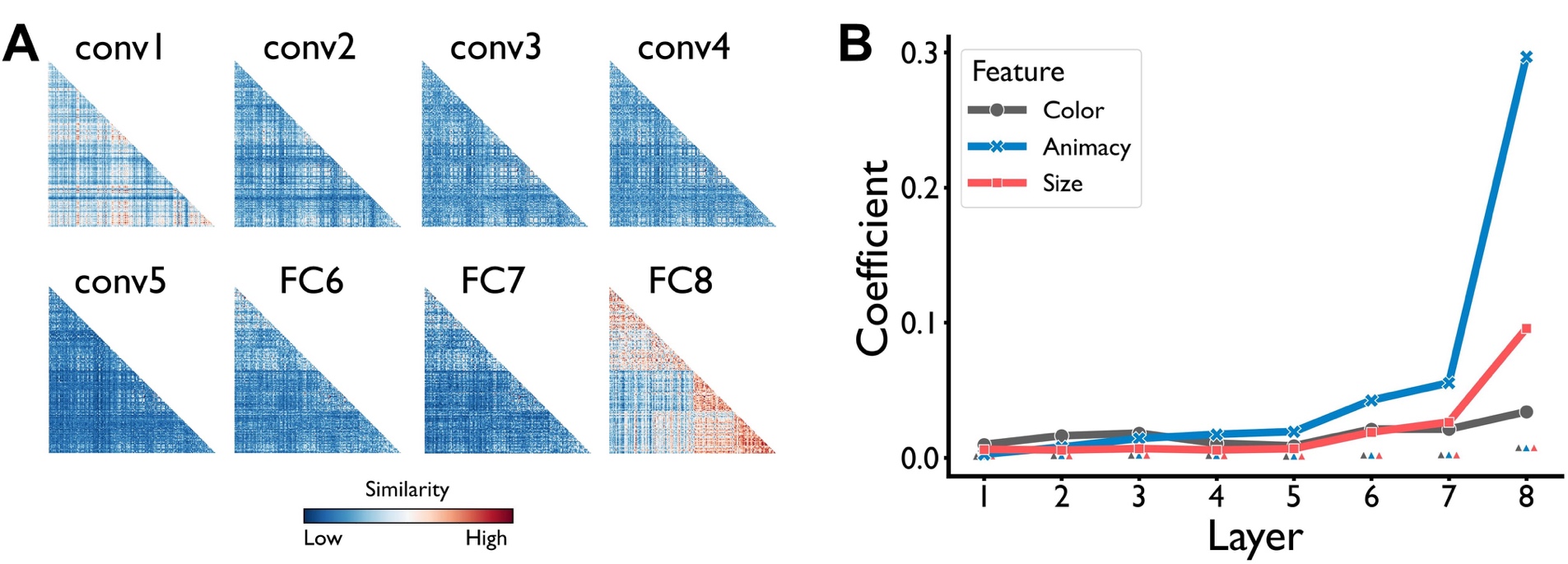

**Figure S2. Representations of three features (color, animacy and size) in the deep neural network AlexNet (stimulus set 2). A. Representational similarity matrix.** The matrix was generated by correlating the activations of the artificial neurons in each DNN layer. **B. Sensitivity of feature information.** The threshold of feature-specific representations in each layer was indicated by the little triangles which were not connected by lines. This figure was generated by the data of the stimulus set 2. conv: convolutional layer; FC: fully connected layer.

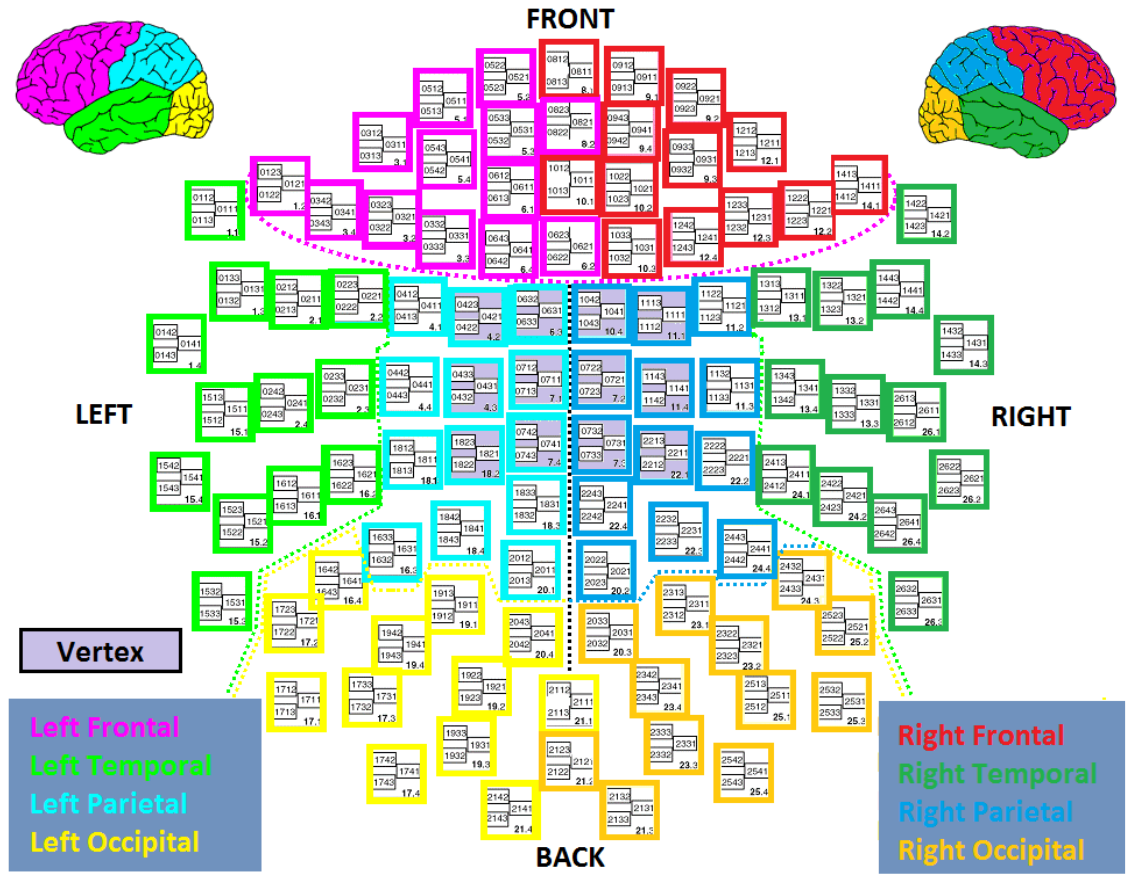

**Figure S3. Elekta Neuromag MEG channel positions.** Channels corresponding to different lobes are color-coded (figure adapted from www.megwiki.org)

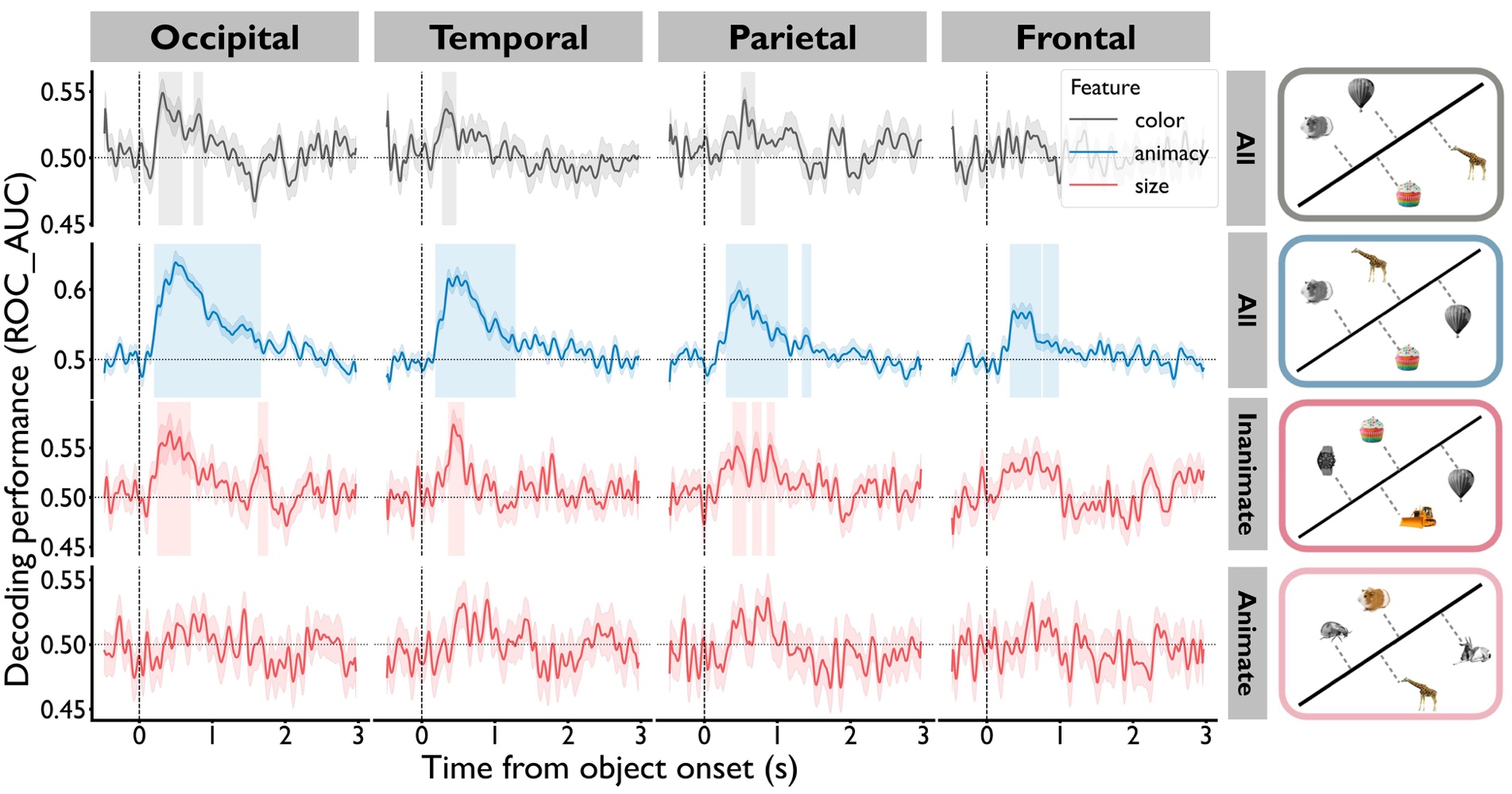

**Figure S4. The decoding performance of three features during the learning stage.** Significant clusters of color, animacy and size features in the occipital, temporal and parietal cortex were found (*P_corrected_* < 0.05). In the frontal cortex, only significant clusters of the animacy feature were found (*P_corrected_* < 0.05). There were no significant clusters of size feature for the animate objects. The shaded areas surrounding the classification performance time courses indicated standard error across participants. The vertical shaded areas marked the significant clusters carrying the feature information.

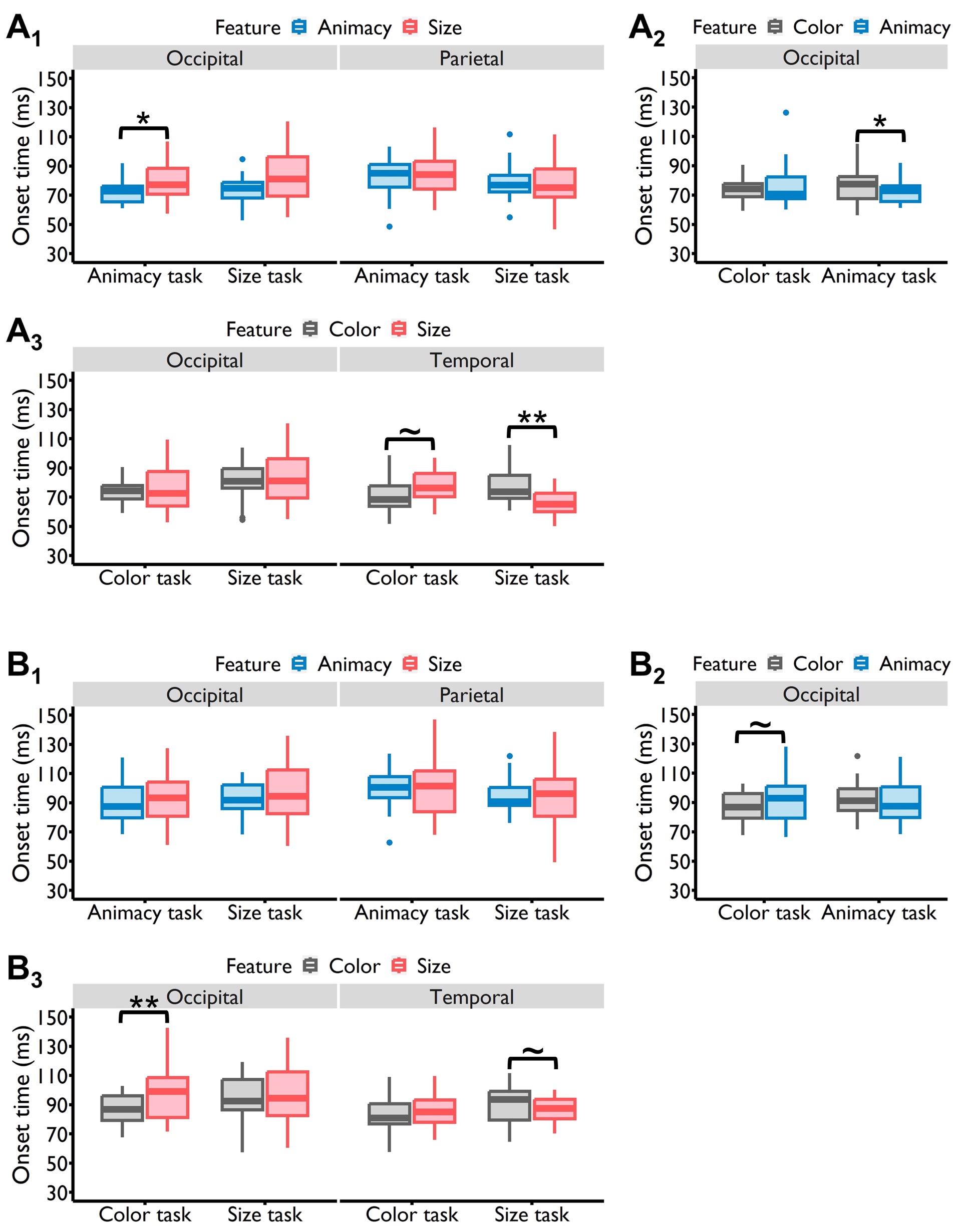

**Figure S5. The modulation effects of selective attention when setting T=30 ms (A) or 40 ms (B)**, i.e., the time window of at least 30 ms or 40 ms in which the d-values of all time points exceeding the corresponding threshold was considered as a significant cluster containing feature information. As shown in the figures, when the minimum duration (T) of a significant cluster increased, the overall onset times of features were delayed accordingly, and the patterns of time lags between different features remained discernible, although some of them became obscured.

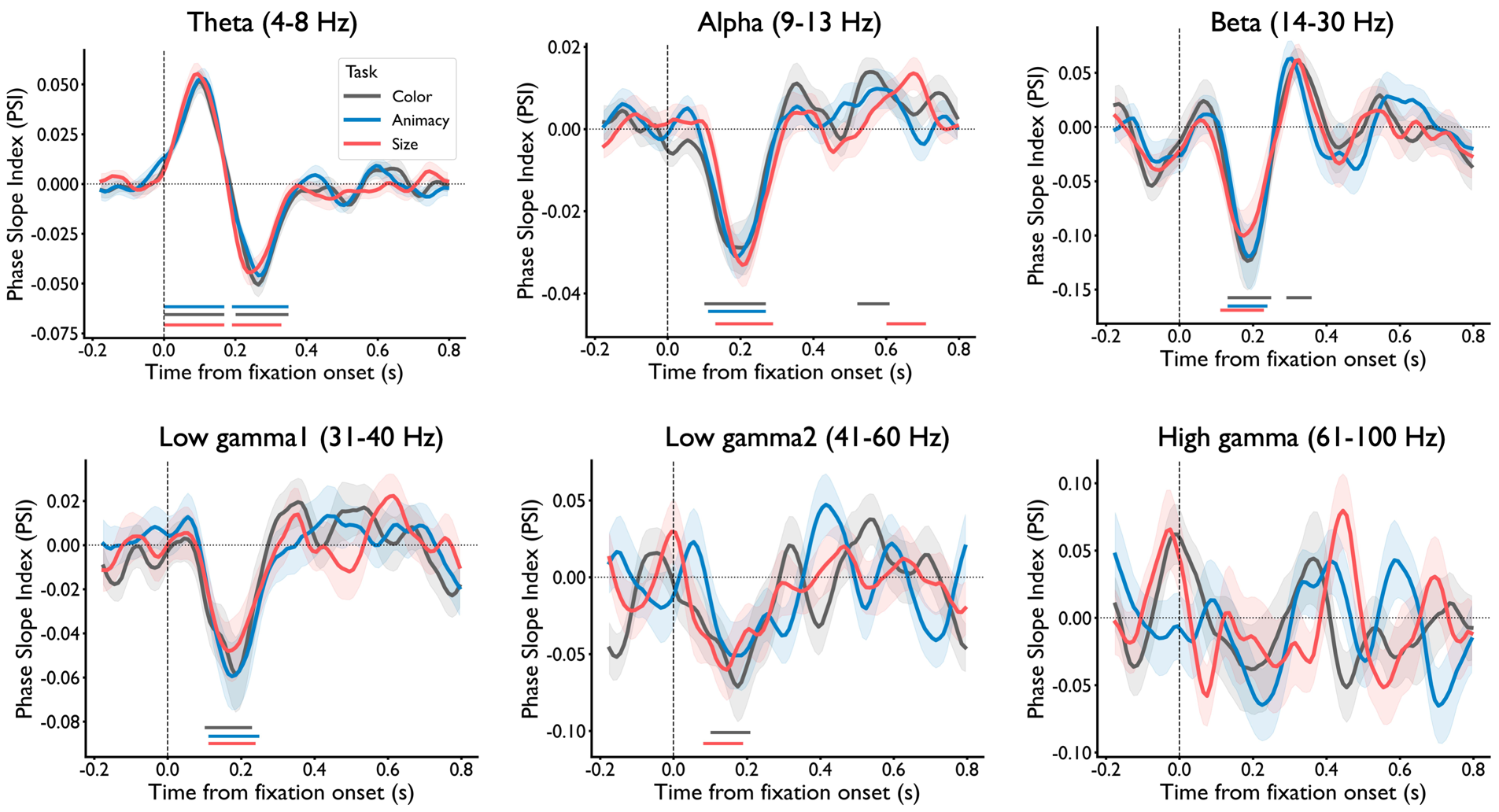

**Figure S6. Phase coupling patterns between the temporal and occipital lobes across different frequency bands.** Only phase coupling in the theta band demonstrated a pattern of initially top-down followed by bottom-up interactions. In contrast, the phase couplings in the alpha, beta, and low gamma bands were predominantly bottom-up, indicating an information flow from the occipital to temporal lobes. No significant phase coupling was observed in the high gamma band.

**Table S1 The coefficients of features for stimulus set 1**

| **Layer** | **Feature** | **Coefficient** | **Threshold** |
| --- | --- | --- | --- |
| 1 | Color | 0.0089 | 0.0014 |
| 2 | Color | 0.0138 | 0.0017 |
| 3 | Color | 0.0156 | 0.0019 |
| 4 | Color | 0.0087 | 0.0016 |
| 5 | Color | 0.0082 | 0.0012 |
| 6 | Color | 0.0190 | 0.0019 |
| 7 | Color | 0.0192 | 0.0023 |
| 8 | Color | 0.0283 | 0.0071 |
| 1 | Animacy | 0.0028 | 0.0012 |
| 2 | Animacy | 0.0075 | 0.0017 |
| 3 | Animacy | 0.0138 | 0.0018 |
| 4 | Animacy | 0.0167 | 0.0015 |
| 5 | Animacy | 0.0187 | 0.0012 |
| 6 | Animacy | 0.0452 | 0.0017 |
| 7 | Animacy | 0.0568 | 0.0023 |
| 8 | Animacy | 0.2878 | 0.0065 |
| 1 | Size | 0.0064 | 0.0012 |
| 2 | Size | 0.0050 | 0.0019 |
| 3 | Size | 0.0064 | 0.0018 |
| 4 | Size | 0.0052 | 0.0015 |
| 5 | Size | 0.0063 | 0.0012 |
| 6 | Size | 0.0186 | 0.0019 |
| 7 | Size | 0.0264 | 0.0025 |
| 8 | Size | 0.0900 | 0.0068 |

**Table S2 The coefficients of features for stimulus set 2**

| **Layer** | **Feature** | **Coefficient** | **Threshold** |
| --- | --- | --- | --- |
| 1 | Color | 0.0099 | 0.0011 |
| 2 | Color | 0.0163 | 0.0016 |
| 3 | Color | 0.0182 | 0.0019 |
| 4 | Color | 0.0107 | 0.0018 |
| 5 | Color | 0.0088 | 0.0012 |
| 6 | Color | 0.0212 | 0.0020 |
| 7 | Color | 0.0212 | 0.0022 |
| 8 | Color | 0.0340 | 0.0076 |
| 1 | Animacy | 0.0026 | 0.0013 |
| 2 | Animacy | 0.0079 | 0.0016 |
| 3 | Animacy | 0.0143 | 0.0020 |
| 4 | Animacy | 0.0174 | 0.0017 |
| 5 | Animacy | 0.0194 | 0.0012 |
| 6 | Animacy | 0.0425 | 0.0019 |
| 7 | Animacy | 0.0554 | 0.0023 |
| 8 | Animacy | 0.2968 | 0.0074 |
| 1 | Size | 0.0062 | 0.0014 |
| 2 | Size | 0.0058 | 0.0015 |
| 3 | Size | 0.0068 | 0.0019 |
| 4 | Size | 0.0058 | 0.0016 |
| 5 | Size | 0.0067 | 0.0012 |
| 6 | Size | 0.0189 | 0.0018 |
| 7 | Size | 0.0264 | 0.0021 |
| 8 | Size | 0.0958 | 0.0075 |
